# Single-Soma Deep RNA Sequencing of Human Dorsal Root Ganglion Neurons Reveals Novel Molecular and Cellular Mechanisms Underlying Somatosensation

**DOI:** 10.1101/2023.03.17.533207

**Authors:** Huasheng Yu, Dmitry Usoskin, Saad S. Nagi, Yizhou Hu, Jussi Kupari, Otmane Bouchatta, Suna Li Cranfill, Mayank Gautam, Yijing Su, You Lu, James Wymer, Max Glanz, Phillip Albrecht, Hongjun Song, Guo-Li Ming, Stephen Prouty, John Seykora, Hao Wu, Minghong Ma, Frank L Rice, Håkan Olausson, Patrik Ernfors, Wenqin Luo

## Abstract

The versatility of somatosensation arises from heterogeneous dorsal root ganglion (DRG) neurons. However, soma transcriptomes of individual human DRG (hDRG) neurons – critical information to decipher their functions – are lacking due to technical difficulties. Here, we developed a novel approach to isolate individual hDRG neuron somas for deep RNA sequencing (RNA-seq). On average, >9,000 unique genes per neuron were detected, and 16 neuronal types were identified. Cross-species analyses revealed remarkable divergence among pain-sensing neurons and the existence of human-specific nociceptor types. Our deep RNA-seq dataset was especially powerful for providing insight into the molecular mechanisms underlying human somatosensation and identifying high potential novel drug targets. Our dataset also guided the selection of molecular markers to visualize different types of human afferents and the discovery of novel functional properties using single-cell *in vivo* electrophysiological recordings. In summary, by employing a novel soma sequencing method, we generated an unprecedented hDRG neuron atlas, providing new insights into human somatosensation, establishing a critical foundation for translational work, and clarifying human species-specific properties.

## Introduction

The somatosensory system conveys senses, such as temperature, touch, vibration, and body position^1^. Primary somatosensory neurons, which convert stimuli to electrical signals, are located in the dorsal root ganglia (DRG) and trigeminal ganglia (TG)^2^. They are greatly heterogeneous, composed of many different molecularly and functionally distinct populations^3^. Somatosensation is fundamental to our daily lives, but becomes a devastating human health problem when malfunctioning, such as during chronic pain and itch. Safe and effective drug options for chronic pain and itch are still limited^4–6^, and the development of novel treatment strategies is greatly needed.

Most of our current knowledge about the mammalian somatosensory system has been obtained from model organisms, mainly rodents. However, the success rate of translating treatment strategies working in model organisms such as rodents to humans has been low^7,8^. There may be multiple reasons for the lack of success, but a noticeable one is species differences of the genetic, molecular and potentially even cellular makeup of somatosensory neurons between rodents and humans. Some significant differences between rodent and human DRG (hDRG) neurons have been noticed in previous studies, with neuropeptides, ion channels, and other markers not always matching between the species^9,10^. For an example, MRGPRX4, a bile acid receptor for human cholestatic itch, does not have a molecular ortholog in mice^11^. In contrast, TGR5, a receptor identified in mouse DRG neurons for bile acid-induced itch^12^, is not expressed in hDRG neurons but instead in the surrounding satellite glial cells^11^. Thus, it is critical to elucidate molecular profiles and cell types of hDRG neurons for understanding human somatosensory mechanisms as well as for translational purposes.

Single-cell RNA sequencing (RNA-seq) is a powerful tool to qualitatively and quantitatively study transcripts of individual cells (soma and/or nuclei)^13^. Based on the transcriptome, including both transcript identities and expression levels, heterogenous cells can be classified into different types^14^. This approach has been successfully used in varying degrees to study DRG and TG neurons in mouse, macaque and other model organisms^15–18^, providing comprehensive molecular and cellular atlases for understanding somatosensory neurons and their differentiation. However, there are some unique technical challenges for conducting single-cell RNA-seq of human DRG or TG neurons: 1) Human tissues are more difficult to obtain compared to model organisms, and the quality of human tissues (RNA integrity) is much more variable; 2) In human DRG/TG tissues, non-neuronal cells, including satellite glial cells, fibroblasts, and other cell types, greatly outnumber the neuronal cells^9,19^; 3) Satellite glial cells tightly wrap around neuronal somas^9,19^, making their physical separation difficult; 4) Human DRG/TG neuronal somas are much larger than those of mouse DRG/TG neuronal somas, so they are prone to damage by enzyme digestion and mechanical forces during single-cell isolation and not compatible with many current sequencing platforms. In addition, transcriptome changes caused by enzymatic and mechanical dissociation may also affect cell clustering and cell type identification^20^. Due to these difficulties, single-nucleus RNA-seq and 10x Visium spatial transcriptomics have been employed for human DRG/TG neurons^21–24^. Despite novel insights from these studies, the quantity of transcripts in the nucleus is much lower than those in the soma, and the nuclear transcripts may not represent the full transcriptome profile of the whole cell^25^, while the 10x Visium spatial transcriptomics lack single-neuron resolution (Fig. 1A). Thus, it is necessary to develop a new method that enables soma RNA-seq of hDRG neurons while preserving the single-cell resolution.

**Figure 1.**
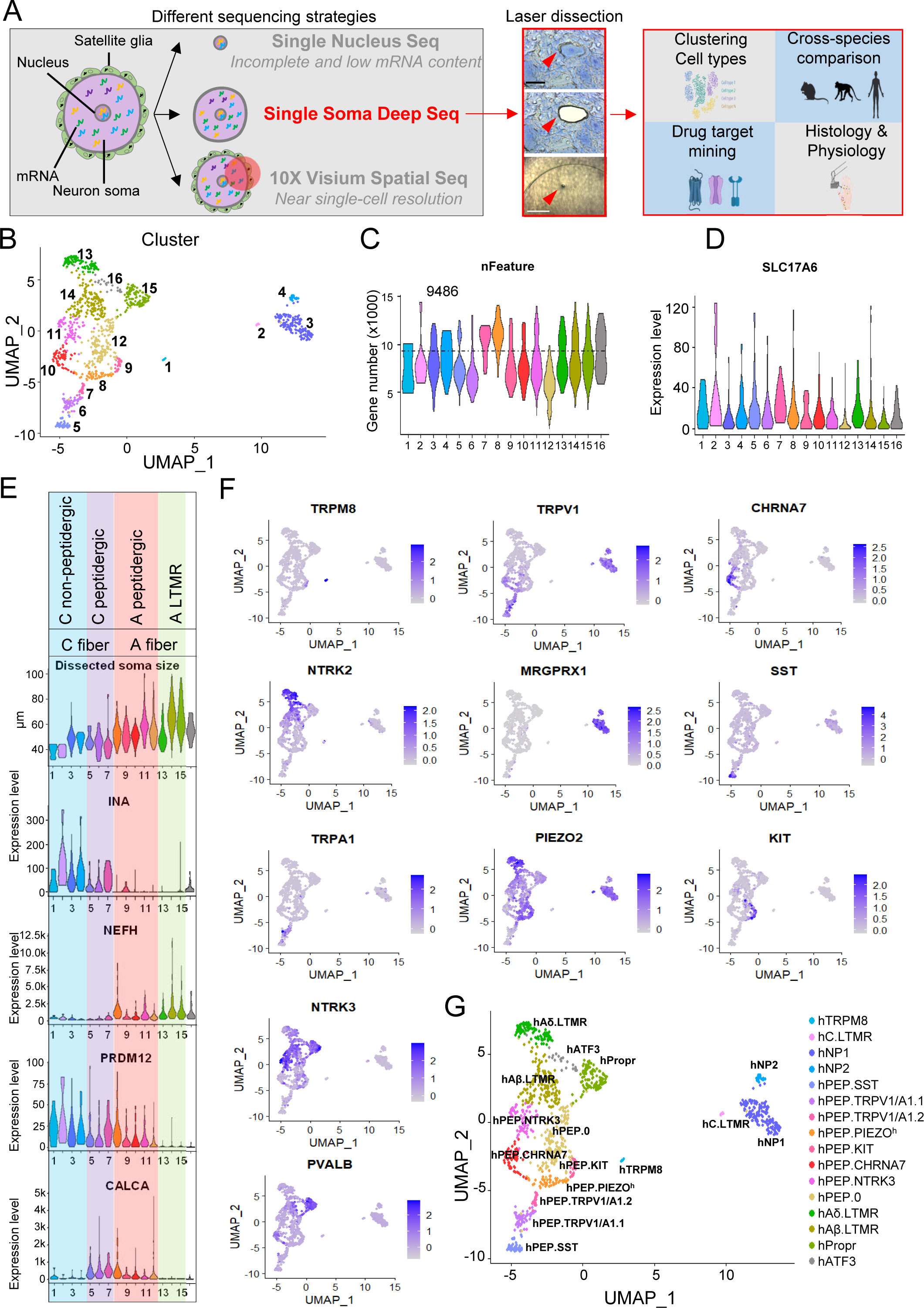
Developing a novel laser capture microdissection (LCM) based approach for single-soma deep RNA-seq of hDRG neurons. (**A**) A diagram shows the overall workflow of this study. (left) A cartoon illustrates features associated with different strategies for single-cell RNA-seq of hDRG neurons. (Middle) One real example of the laser dissection of a hDRG neuron soma is shown. (Right) Analyses and experiments conducted in this study are summarized. Scale bar, 50 μm (cell) and 500 μm (cap) (**B**) UMAP plot showing the clusters of 1066 hDRG neurons. (**C-D**) Violin plots showing total number of detected genes (C) and the expression of neuronal marker *SLC17A6* (D). (**E**) The grouping clusters based on the soma size, and the expression of *INA*, *NEFH*, *PRDM12*, and *CALCA*. (**F**) UMAPs showing some canonical marker gene expression in each cluster. (**G**) UMAP plot with names of each cluster.

To meet this challenge, we developed a novel approach by combining laser capture microdissection (LCM) for isolating individual neuronal somas and the Smart-seq2^18^ for generating full length RNA-seq libraries (Fig. 1A). We sequenced 1066 hDRG neurons with minimum satellite glial cell contamination from six lower thoracic and lumbar DRGs of three donors, detecting an average of >9,000 unique genes per neuron (∼ 3-5 times or more than the previous single-nucleus RNA-seq results) and identifying 16 molecularly distinctive neuron types. Crossspecies analysis revealed both similarities but considerable differences among human, macaque, and mouse DRG neurons. In addition, we uncovered a set of novel marker genes that can help to distinguish different types of sensory neurons and afferents in hDRG and skin tissues. Based on the molecular profiles, we also predicted novel response properties of human sensory afferents, which were tested and confirmed by single-cell *in vivo* recordings. These results support a close relationship between the molecular profiles uncovered by single-soma RNA-seq, histology, and functional properties of human sensory afferents, highlighting the precision and unique utility of this dataset in guiding functional studies of human somatosensory neurons. In short, we have established a novel approach to conduct single-soma deep RNA-seq of hDRG neurons, which revealed previously unknown neuronal types and functional properties. Given the high number of unique transcripts recovered from each neuron, our dataset is especially powerful for molecular discovery, such as identifying potential high value novel drug targets. We believe that this high-fidelity single-soma RNA-seq dataset will serve as a ground reference for homogenizing RNA-seq data of human DRG/TG neurons using different approaches and for translating animal studies into therapeutic applications.

## Results

### Development of an LCM-based approach for conducting single-soma deep RNA-seq of human DRG neurons

Adult human DRG neurons present considerable technical challenges for high-quality single-cell RNA-seq study. To overcome these hurdles, we developed a novel method that utilizes the laser capture microdissection to isolate individual neuronal soma and combine with Smart-seq2 full length RNA-seq library generation method for single-soma high-depth transcriptomic analysis (Fig. 1A). Six hDRGs at the low thoracic (T11-T12) and lumbar (L2-L5) levels from three postmortem donors (donor information and screening criteria are summarized in the supplementary tables 1 & 2) were procured through NDRI (National Disease Research Interchange). DRGs were extracted and immediately frozen in OCT (within ∼ 6 hours after death) at the NDRI procurement sites to minimize RNA degradation. Fresh frozen DRGs were cryo-sectioned, mounted onto LCM slides, and briefly stained with a HistoGene^TM^ dye for cell visualization (Fig. S1A-C). Individual neuronal somas were discernable under microscope and dissected by a laser (Fig. 1A). Each detached soma dropped into a tube cap containing cell lysis buffer for library preparation (Fig. 1A). Dissected neuronal somas exhibited a similar size distribution as the whole DRG neuron population (Fig. S1D-E), suggesting no obvious sampling bias. Sequencing libraries were generated following the Smart-seq2 protocol^26^. In total, 1136 neuronal somas were dissected and passed through the final quality control for sequencing. During preliminary bioinformatic analysis, 70 samples with obvious glial cell contamination (dominant expression of APEO, FABP7, and other glia cell markers) were removed, and the remaining 1066 neurons were used for analysis in this study. 16 transcriptomic clusters of hDRG neurons were identified by Seurat^27^ (Fig. 1B), with an average of 9486 genes detected per cell (Fig. 1C). No obvious batch effects or do-nor differences were observed in the clusters (Fig. S2A-E). All these cells expressed high levels of known peripheral sensory neuronal markers, *SLC17A6* (*VGLUT2*), *SYP* (Synaptophysin), and *UCHL1* (*PGP9.5*) (Fig. 1D, S2F-G).

### Anatomical and molecular features of hDRG neuron clusters

Mammalian DRG neurons have some well-known physiological, anatomical, and molecular features. DRG somatosensory afferents can be identified as A- or C- afferent fibers according to axon conduction velocities^28^. A-fiber DRG neurons usually have large-diameter somas and myelinated axons, while C-fiber DRG neurons have small-diameter somas and non-myelinated axons. The A- and C- afferents can be further divided into peptidergic and non-peptidergic types, based on the expression of one neuropeptide, calcitonin related polypeptide alpha (*CALCA(*CGRP))^29^. To determine the broad types of the 16 hDRG neuron clusters, we analyzed their soma sizes and expression of some key molecular markers (Fig. 1E). Clusters 1-16 were arranged from small to large soma sizes. In addition, expression of neurofilament intermediate filament (*INA*), which was enriched in small-diameter DRG neurons, and heavy chain (*NEFH*), which was highly expressed in large-diameter DRG neurons, showed a clear complementary pattern. The combined morphological and molecular features suggested that clusters 1-7 were likely to be unmyelinated, small diameter C-fiber DRG neurons, whereas clusters 8-16 were myelinated, large diameter A-fiber DRG neurons (Fig. 1E). The two groups could be further divided based on the expression of *CALCA* (clusters 5-12). Moreover, the PR domain zinc finger protein 12 (*PRDM12*), a transcriptional regulator critical for development of human pain-sensing afferents (nociceptors) and C fibers^30^, was expressed in clusters 1-12, further distinguishing between C- and A-fiber nociceptors versus A-fiber low-threshold mechanoreceptors (Fig. 1E). Taken all information into consideration, clusters 1-4 were classified as non-peptidergic C-fibers, clusters 5-7 as peptidergic C-fibers, clusters 8-12 as peptidergic A-fibers, clusters 13-15 as low-threshold mechanoreceptors A-fibers (A-LTMRs), and cluster 16 as an unknown A-fiber type (Fig. 1E).

Based on the expression profiles of top molecular markers, which had either unique expression pattern or well-known functions (Fig. 1F, S2H), we named these 16 hDRG neuron clusters using a nomenclature system with the following format (Fig. 1G): 1) The “h” at the beginning of each cluster name indicated “human”; 2) mouse nomenclature was used for conserved subtypes (i.e. most non-peptidergic neurons and A-LTMRs); 3) human peptidergic neuron types were designated as hPEP.(marker gene). Briefly, for non-peptidergic C-fiber neurons, cluster 1 was named hTRPM8, cluster 2 was C-fiber low-threshold mechanoreceptors (hC.LTMR), cluster 3 was type I non-peptidergic nociceptors (hNP1), and cluster 4 was type II non-peptidergic nociceptor (hNP2). For peptidergic C-fiber neurons, cluster 5 was named hPEP.SST, cluster 6 hPEP.TRPV1/A1.1, and cluster 7 hPEP.TRPV1/A1.2. For peptidergic A-fiber neurons, cluster 8 was named hPEP.PIEZO^h^ (the superscript ‘h’ means ‘high’), cluster 9 hPEP.KIT, cluster 10 hPEP.CHRNA7, cluster 11 hPEP.NTRK3, and cluster 12 hPEP.0 (no distinctive molecular marker). For A-LTMRs, cluster 13 was named Aδ low-threshold mechanoreceptors (hAδ.LTMR), cluster 14 Aβ low-threshold mechanoreceptors (hAβ.LTMR), and cluster 15 proprioceptors (hPropr). Cluster 16 was named hATF3.

To validate the clusters identified by Seurat analysis, we independently analyzed our data using the graph-based clustering Conos^31^ package. Cluster structure revealed by Conos analysis reproduced the Seurat results (Fig. S3A). As a third method to validate identified clusters, we employed a neural network-based probabilistic scoring module^15,18^ that learned human cell type features based on their molecular profiles (Fig. S3B-D). Namely, the accuracy score of our Seurat clustering assignment by the learning module was near 90% (Fig. S3B), which meant that most cells were accurately assigned to their corresponding clusters, confirming the accuracy of clustering (Fig. S3C). Moreover, the cell type consistency was validated by probabilistic similarity (Fig. S3D). Thus, three independent analysis methods confirmed the robustness of the clustering and strongly supported the cell type assignment.

Using Conos we also performed co-clustering and label propagation with a recently published single-nucleus RNA-seq dataset of hDRG neurons by Nguyen et al^23^ (Fig. S4). This analysis showed that while some clusters displayed a one-to-one match between the two datasets, such as hTRPM8, hPEP.SST, hPEP.KIT, and hAd.LTMR, most did not have one-to-one match. This mismatch could be caused by biological variations such as nucleus vs cytoplasm RNA species and quantity^25^ and technical differences (the increased resolution obtained by deep sequencing in the present study or variability caused by the different technology platforms^26,32^). This mismatch also indicated that our dataset generated novel molecular profiles and cell types different from the single-nucleus RNA-seq results.

### Cross-species comparison of DRG neuron types

Comparison of hDRG neuron types to those in model organisms helps to uncover the evolutional conservation and divergence of DRG neuron populations, provides clues about functions of hDRG neuron populations, and identifies potential species-specific hDRG neuron populations. Here we performed a cross-species comparison between our human dataset, a mouse dataset (Sharma)^16^, and a macaque dataset (Kupari, SmartSeq2 dataset)^18^. To identify corresponding clusters between human and previously published mouse and macaque datasets, we used three different strategies: Conos pairwise co-clustering followed by label propagation (Fig. 2A-B & S5A-B), probabilistic neural network learning (Fig. S5C-D), and machine-learning based hierarchical clustering of an integrated dataset of human and macaque (Fig. S5E). For characteristics and details of different approaches, see Method section. In all these analyses, human non-PEP DRG neuron cell types showed high correlation to those of mouse and macaque, including hC.LTMR, hNP1, hNP2, hPEP.SST (called NP3 in mouse and monkey), hTRPM8, hAδ.LTMR, hAβ.LTMR, and hProprioceptor neurons (Fig. 2A-B & S5). hPEP.TRPV1/A1.1 and hPEP.TRPV1/A1.2 corresponded to macaque PEP1 (Usoskin nomenclature) and mouse subclass PEP1.4/CGRP-ε. This conclusion was supported by both Conos label propagation analysis (Fig. 2A-B) and probabilistic neural network analysis (Fig. S5C-D), suggesting that these clusters to represent C-fiber thermoreceptors and nociceptors. Notably, four types of mouse C-fiber PEP (CGRP) nociceptors have been evidenced^18,33^ (nomenclature from Emory & Ernfors/Sharma), but our analysis indicated that mouse clusters PEP1.1/CGRP-α, PEP1.2/CGRP-β, and PEP1.3/CGRP-γ might not be evolutionarily conserved, as they did not have equivalent types in the human datasets (Fig. 2A & S5C). hPEP.CHRNA7 showed high correlation to mouse PEP2/CGRP-ζ and macaque PEP2, while hPEP.KIT corresponded to mouse PEP3/CGRP-η and macaque PEP3, suggesting these clusters functionally belong to A-fiber nociceptors (Fig. 2A-B & S5C-D). Interestingly, hPEP.NTRK3, hPEP.PIEZO^h^, and hPEP.0, which made about half of human PEP nociceptors that we sequenced, did not have a one-to-one corresponding cell types in either mouse or macaque (Fig. 2A-B & S5C-D). Since macaque PEP2 cluster in the high-quality Smart-Seg2 macaque dataset contained only three neurons, we focused our interpretation on analyses on mouse homologs for this cell type. hPEP.NTRK3 showed the greatest similarity to mouse PEP2 (CGRP-ζ) by Conos propagation analysis (Fig. 2A). Probabilistic neural network learning revealed similar scores to both mouse PEP2 (CGRP-ζ) and PEP3 (CGRP-η) (Fig. S5C). Overall, it therefore seems that hPEP.NTRK3 represents a convergent mouse PEP2/3-like cell type. hPEP.PIEZO^h^ showed some similarity to mouse PEP3 (CGRP-η) and macaque PEP1 and PEP3 in co-clustering (Fig. 2A-B). Neural network learning, and hierarchical clustering, indicating this cell type to represent an A-fiber mechanosensory nociceptor (Fig. S5C-E). Thus, hPEP.NTRK3 and hPEP.PIEZO^h^ represents diverged PEP2- and PEP3-like A-fiber nociceptors, which likely have emerged as human species-specific sensory neuron types. hPEP.0, a type of human PEP A-afferents, showed no similarity to any mouse DRG neurons but some relationship to macaque PEP1 and PEP3 (Fig. 2A-B), suggesting that they might be primate specific PEP nociceptors. A schematic overview of our conclusions based on the above analyses regarding the cell-type homologs across mouse-macaque-human is illustrated in Fig. 2C.

**Fig. 2.**
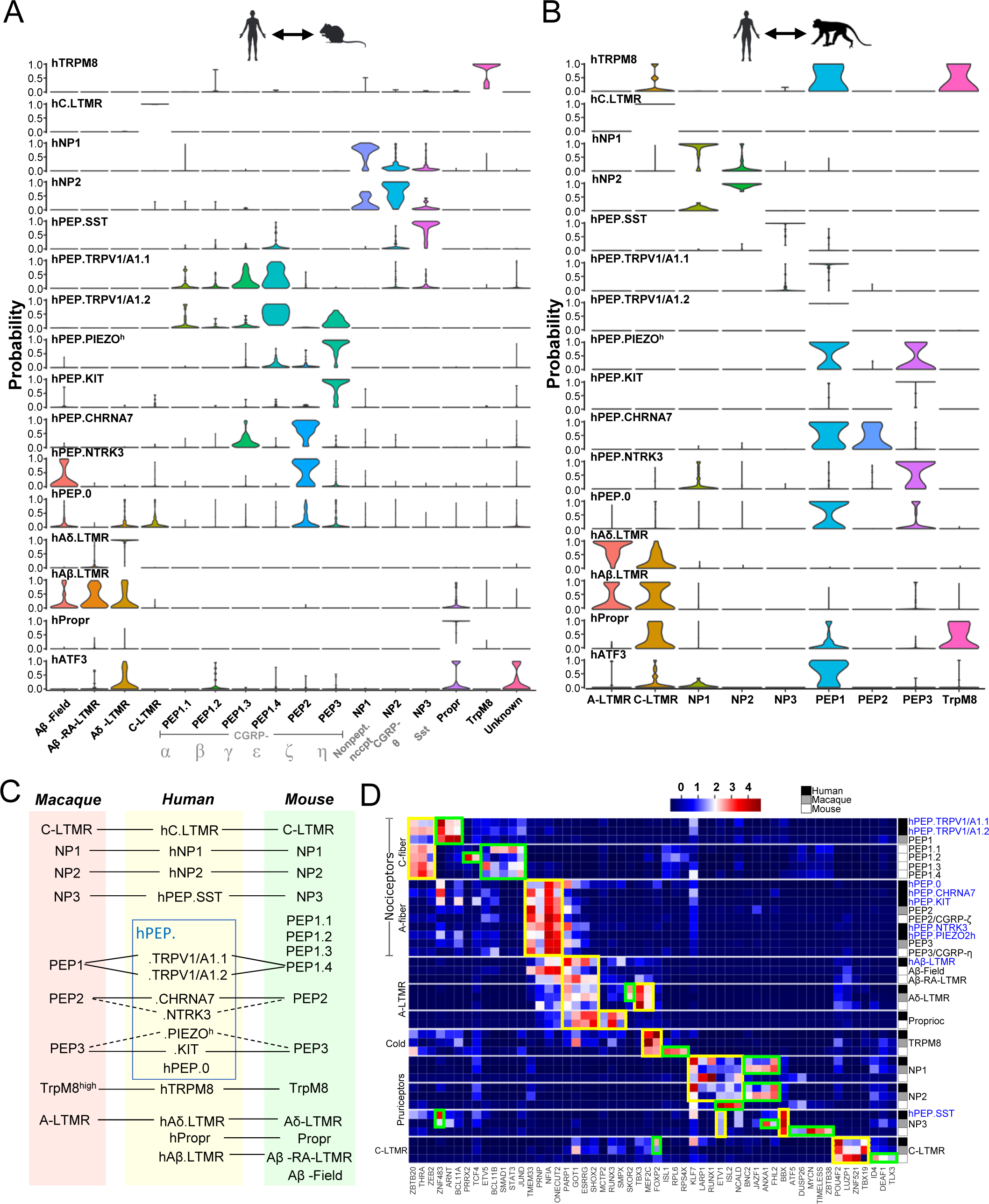
Cross-species analysis of DRG neurons in human, macaque, and mouse. (**A, B**) Conos label propagation from mouse (Sharma) (A, combined Sharma & Usoskin nomenclature) and macaque (B) to hDRG neuron clusters showing the cell type correlation. For UMAPs for correspondent co-integration, from which these results were inferred, see Fig. S5A-B. **(C**) Summary of correspondence of DRG neuron clusters among three species. Solid lines depict clear match, and dashed lines represent partial similarity. (**D**) Heatmap visualization of cross-species-conserved and species-specific transcription factor associated gene patterns across mouse, macaque, and human. Species are color coded in the right column. Yellow boxes, conserved, green boxes, species-specific gene regulatory networks.

Transcription factors play critical role in DRG neuron development and differentiation^34^. Thus, we performed a transcription factor associated gene regulatory network analysis (TF-GRNs) using machine learning to identify shared and species-specific TF-GRNs contributing to the similarities and differences between sensory neuron subtypes and species (Fig. 2D). Evolutionarily conserved TF-GRNs defining C-fiber nociceptors, A-fiber nociceptors, A-LTMRs, TRPM8, C-fiber pruriceptors/nociceptors (hNP1, hNP2, hPEP.SST), and C-LTMRs were observed (yellow boxes in Fig. 2D) as well as species-specific networks, such as for C-fiber nociceptors, hTRPM8, hNP1, hNP2, hPEP.SST and hC.LTMR (green boxes in Fig. 2D). Among cross-species conserved transcriptional regulators, some were previously known to drive sensory neuron diversification in mouse, including *ZEB2* in C-fiber nociceptors^35^, *SHOX2* in A-LTMRs^36,37^, *RUNX3* in proprioceptors^38^, *FOXP2* in TRPM8, *RUNX1* in NP1^39^, *ZNF52* and *POU4F2* in C-LTMRs^16,40^. These transcription factors may contribute to the formation of different DRG neuron cell types and regulate the conserved and species-specific gene expression in each cell population.

### Similarities and differences of top marker genes across species

To reveal molecular differences among the corresponding cell types in human, macaque (Kupari)^18^, and mouse (Sharma)^16^, we selected the top 10 marker genes from each hDRG neuron population and compared them across species (Fig. S6). In general, the expression patterns of these genes were more similar between human and macaque than between human and mouse, reflecting the evolutionary distances between rodent, non-human primate, and human. Some genes were expressed in the corresponding populations across all three species. For example, *TRPM8* was expressed in C-fiber cold-sensing neurons, and *IL31RA* was expressed in nociceptive/pruriceptive population (NP3) as well as its corresponding human population hPEP.SST. Some genes, such as *KCNV1*, a voltage-gated potassium channel, were specific for primate Aδ.LTMR but had low or no expression in the corresponding mouse DRG neurons (Fig. S6). Moreover, some marker genes were specific only for hDRG neuron types. For example, Mechanosensory Transduction Mediator Homolog (*STUM*) was specifically expressed in hTRPM8, and calsequestrin 2 (*CASQ2*) was specifically expressed in hC.LTMR (Fig. S6). Genes specifically enriched in hDRG neurons may confer unique physiological and functional properties of the human somatosensory system. The observed differences from the top 10 marker genes represent only the iceberg tip of the overall molecular differences between human and model organism DRG neurons, highlighting the necessity of validating molecular targets in hDRG neurons for translational studies.

### Molecular marker expression and validation of C-fiber pruriceptors, thermoreceptors and nociceptors

Based on the sequencing results, we established specific marker genes or their combination to identify each type of hDRG neurons (Fig. S7A) and validated their expression using RNASCOPE multiplex fluorescent *in situ* hybridization (FISH) (Fig. S8). We also deduced potential functions of each hDRG neuron type based on the cross-species cell type comparison and expression of molecules with known sensory functions.

hNP1 and hNP2 exclusively expressed *MRGPRX1* (Fig. 3A), a Mas-related GPCR family member that could be activated by various pruritogens^41^. Similarly, primate specific bile acid receptor *MRGPRX4* for mediating human cholestatic itch^11,42^ were detected by our single-soma dataset. Enrichment of *MRGPRX4* and *MRGPRX3* (an orphan GPCR in the same family) in hNP2 but not hNP1 helped to separate these two clusters (Fig. S7A). Other itch-related receptors, such as *HRH1* and *IL31RA* (Fig. 3A-B), were also expressed in both hNP1 and hNP2, suggesting that these two populations function to detect various pruritogens and transmit the sensation of itch.

**Figure 3.**
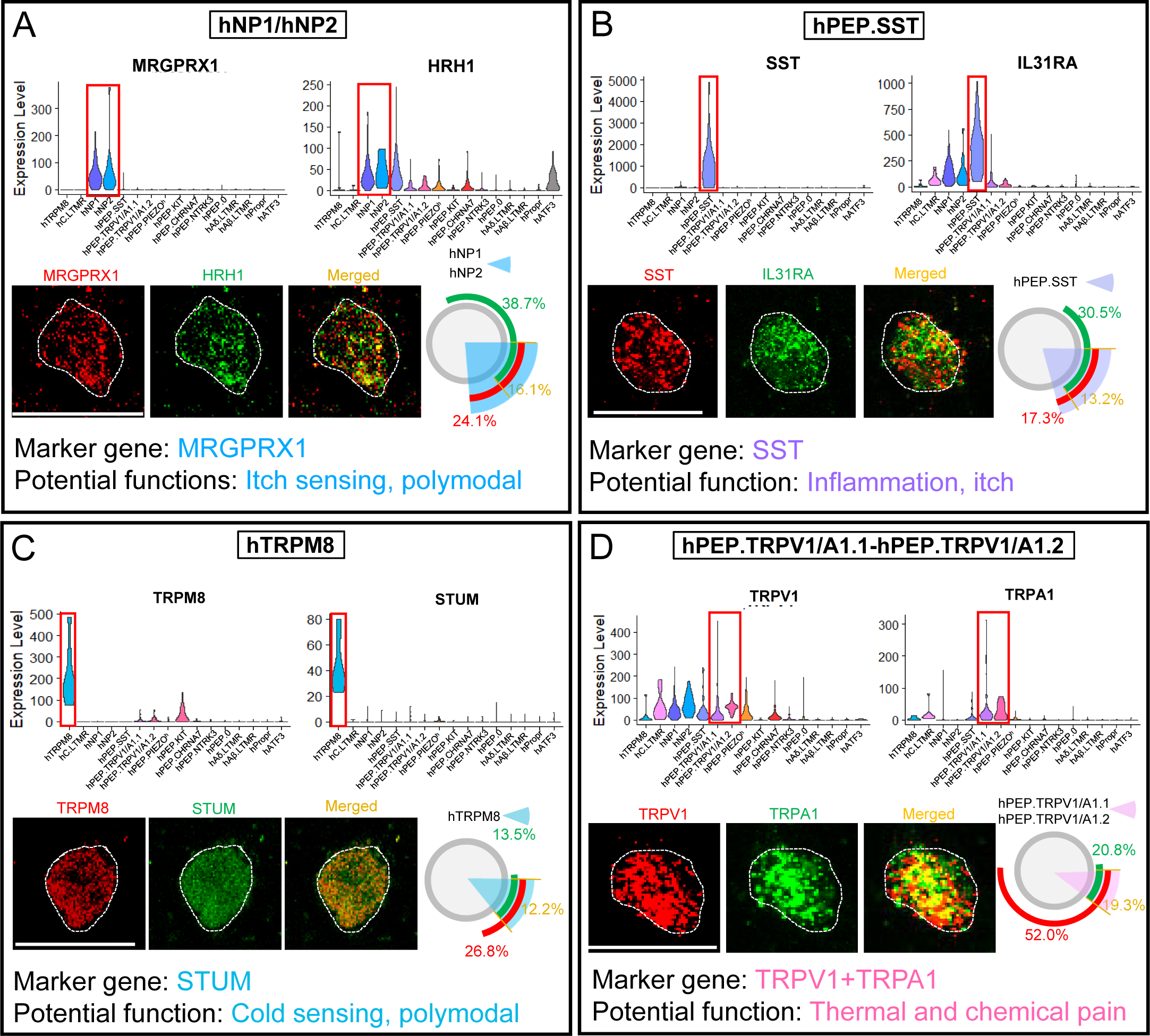
Marker gene expression and validation in C-fiber pruriceptors, thermoreceptors, and nociceptors. (**A-D**) Marker genes for specific labeling of each cluster and validation by multiplex FISH for hNP1 and hNP2 (A), hPEP.SST(B), hTRPM8 (C), and hPEP.TRPV1/A1.1 and hPEP.TRPV1/A1.2 (D). The fluorescent images show detected transcripts in one example hDRG neuron (cell body outlined by the white dashed line. Circle charts next to images show quantification: the arcs indicate the percentage of neurons positive for the given marker gene over all sampled DRG neurons. The sector shaded areas indicate the approximate percentage of each cell type over the total quantified hDRG neurons. N=2, A (199 neurons total), B (220 neurons total), C (156 neurons total), D (202 neurons total). Scale bar, 50 μm.

The co-expression of *MRGPRX1* and *HRH1* was validated by multiplex FISH (Fig. 3A). *PIEZO2* was expressed at a higher level in hNP1 than in hNP2 neurons (Fig. S7B), suggesting that hNP1 neurons could be more mechanosensitive. Consistent with this molecular feature, recordings from human afferents have revealed that some human histaminergic itch-sensing fibers are mechanosensitive^43^. In mice, NP1 and NP2 afferents are well-characterized by specific expression of two different *Mrgpr* members: NP1 neurons (∼20% of total DRG neurons) highly express *Mrgprd*^44^, while NP2 neurons (∼5% of total DRG neurons) express *MrgprA3*^15^. In humans, however, *MRGPRD* was only expressed in a few NP1 neurons (Fig. S9A), so it was less useful for marking human NP1 population, and *MRGPRA3* gene does not have an orthologue in the human genome. In short, NP1 and NP2 populations likely have conserved physiological functions in mediating itch sensation in both mice and humans. However, some key molecular receptors for detecting pruritogens are different between the species, which may reflect evolutionary adaptation to distinctive pruritogens encountered by human and mice in their living environments.

The hPEP.SST population displayed specific expression of the neuropeptide, somatostatin (*SST*), and an enriched expression of *GFRA3,* a co-receptor of tyrosine kinase RET (Fig. 3B, S7A). We also found another neuropeptide cholecystokinin (*CCK*)^45^ enriched in this population (Fig. S7A). The hPEP.SST cluster corresponded to mouse and macaque NP3 population (Fig. 2), which are also marked by the expression of *SST*^15,16^. Given the previously established role of mouse NP3 neurons in itch sensation^46^ and the high expression of itch-sensing receptors, such as *HRH1*, *IL31RA* (Fig. 3A-B), hPEP.SST afferents could also mediate itch sensation, especially under inflammatory conditions^15^. Neither *PIEZO1* nor *PIEZO2* (Fig. S9A) was detected in hPEP.SST neurons, indicating that these afferents might not be mechanosensitive. Indeed, some human histaminergic itch-sensing fibers are insensitive to mechanical forces^43^. A human feature of hPEP.SST neurons was the co-expression of the peptidergic neuron marker, CGRP, which was barely detected in the corresponding mouse NP3 neurons (Fig. S9B)^16,18^.

The hTRPM8 population was distinguished from other cell types by the specific expression of a novel molecular marker, *STUM,* and high-level expression of *TRPM8* (Fig. 3C). Almost all *STUM*^+^ neurons expressed *TRPM8,* which was validated by multiplex FISH (Fig. 3C). Since *TRPM8* is a receptor for cold temperature and cooling chemicals (such as menthol)^47–50^, the hTRPM8 population likely functions as cold- and menthol-sensing afferents. Notably, this newly identified maker gene *STUM* was not detected in mouse *TRPM8^+^*neurons^16,33^. In macaque, *STUM* was more broadly expressed in several clusters (Fig. S9C)^18^. Nevertheless, other molecular markers, such as *FOXP2* and *GPR26*, were shared among mouse^16,33^, macaque^18^, and human TRPM8 cold-sensing neurons (Fig. S9A-C). Intriguingly, some hTRPM8 neurons also expressed a low level of heat-sensing receptor *TRPV1* (Fig. 3D), suggesting that these neurons might also be activated by heat stimuli. Consistently, human physiological recordings have identified cold-sensitive C-fibers that also respond to heat^51^. Thus, some of the neurons in the hTRPM8 population are likely to be polymodal cold-sensing afferents.

Two peptidergic C-fiber clusters displayed overlapping high expression of *TRPV1* and *TRPA1* (Fig. 3D), which were therefore named as hPEP.TRPV1/A1.1 and hPEP.TRPV1/A1.2. Since *TRPV1* is activated by noxious heat and capsaicin, and *TRPA1* can be activated by noxious cold and various noxious chemicals^52,53^, they are likely to be C-fiber peptidergic thermoreceptors and nociceptors, sensing noxious thermal and chemical stimuli and transmitting pain signals. From the cross-species comparison, hPEP.TRPV1/A1.1 and hPEP.TRPV1/A1.2 were mostly similar to macaque PEP1 and mouse subclass PEP1.4/CGRP-ε (Fig. 2A-C). The exact functional differences between these two quite similar populations are yet to be established, but it is tempting to hypothesize that hPEP.TRPV1/A1.1 innervates the skin while hPEP.TRPV1/A1.2 innervates deep tissues and visceral organs, because CGRP^+^/TRPV1^+^ afferents were observed in both the human skin and deep tissues^54^. In contrast to the hPEP.TRPV1/A1.1 population, hPEP.TRPV1/A1.2 neurons expressed PROKR2, and higher level of PTGER3, and prostaglandin I2 receptor (PTGIR) (Fig. S7E & S9A). These molecular markers have been found to be enriched in mouse viscera-innervating DRG neurons^55,56^.

### Molecular marker expression and validation of A-fiber peptidergic nociceptors

Five clusters of peptidergic A-fiber nociceptors were identified in our dataset. When compared to mouse and macaque peptidergic populations, some clusters showed the greatest divergence (Fig. 2A-C), indicating that they might contain human specific sensory neuron types.

The hPEP.PIEZO^h^ cluster was named because they expressed the highest number of *PIEZO2* transcripts among all PEP clusters (Fig. 4A), an expression level comparable to the most mechanosensitive afferents, hC.LTMR and hAδ.LTMR. The hPEP.PIEZO^h^ neurons also expressed a relatively high levels of *PIEZO1* transcripts, though the overall expression of *PIEZO1* in hDRG neurons was low (Fig. S9A). This cluster could be identified by its high expression of *PTGER3* but not *TRPA1* (Fig. 4A & 3D). No mouse DRG neuron population was highly matched for the hPEP.PIEZO^h^ cluster. Nevertheless, the expression of a few unique marker genes in this cluster could provide clues into its functions. The adrenoreceptor, *ADRA2C*, a specific molecular marker found in human sensory fibers innervating blood vessels^57–60^, was mainly detected in this cluster (Fig. S9A). In addition, the hPEP.PIEZO^h^ population expressed *GPR68*, a membrane receptor sensing flow and shear forces in the vascular endothelia cells^61^ (Fig. S9A). Given the well-established functions of *PIEZO1* and *PIEZO2* in mediating neuronal sensing of blood pressure and the baroreceptor reflex^62^ and the expression of known function of *ADRA2C* and *GPR68*, we proposed that some hPEP.PIEZO^h^ afferents innervate blood vessels and sense the blood pressure or flow. This cluster also expressed a high level of *PTGIR* (Fig. S9A). Mouse *PTGIR*^+^ DRG neurons innervate the bladder^55^, and *PIEZO2* expressed in these neurons is required for sensing the bladder pressure to coordinate urination^63^. Thus, some hPEP.PIEZO^h^ afferents might also innervate bladder and play a similar role. Take all into consideration, hPEP.PIEZO^h^ neurons might function in sensing mechanical forces from blood vessels and internal organs. Given that there was not a clear mouse DRG CGRP^+^ population with high *PIEZO2* expression from single-cell RNA-seq datasets, we speculated that hPEP.PIEZO^h^ neurons are either human specific or greatly expanded in humans. A fundamental difference between human and mouse is their body sizes (humans are more than 2000 times larger than mice^64^). This addition or expansion of hPEP.PIEZO^h^ neurons is likely to meet the challenge of mediating sensation from internal organs and blood vessels in much larger human bodies.

**Figure 4.**
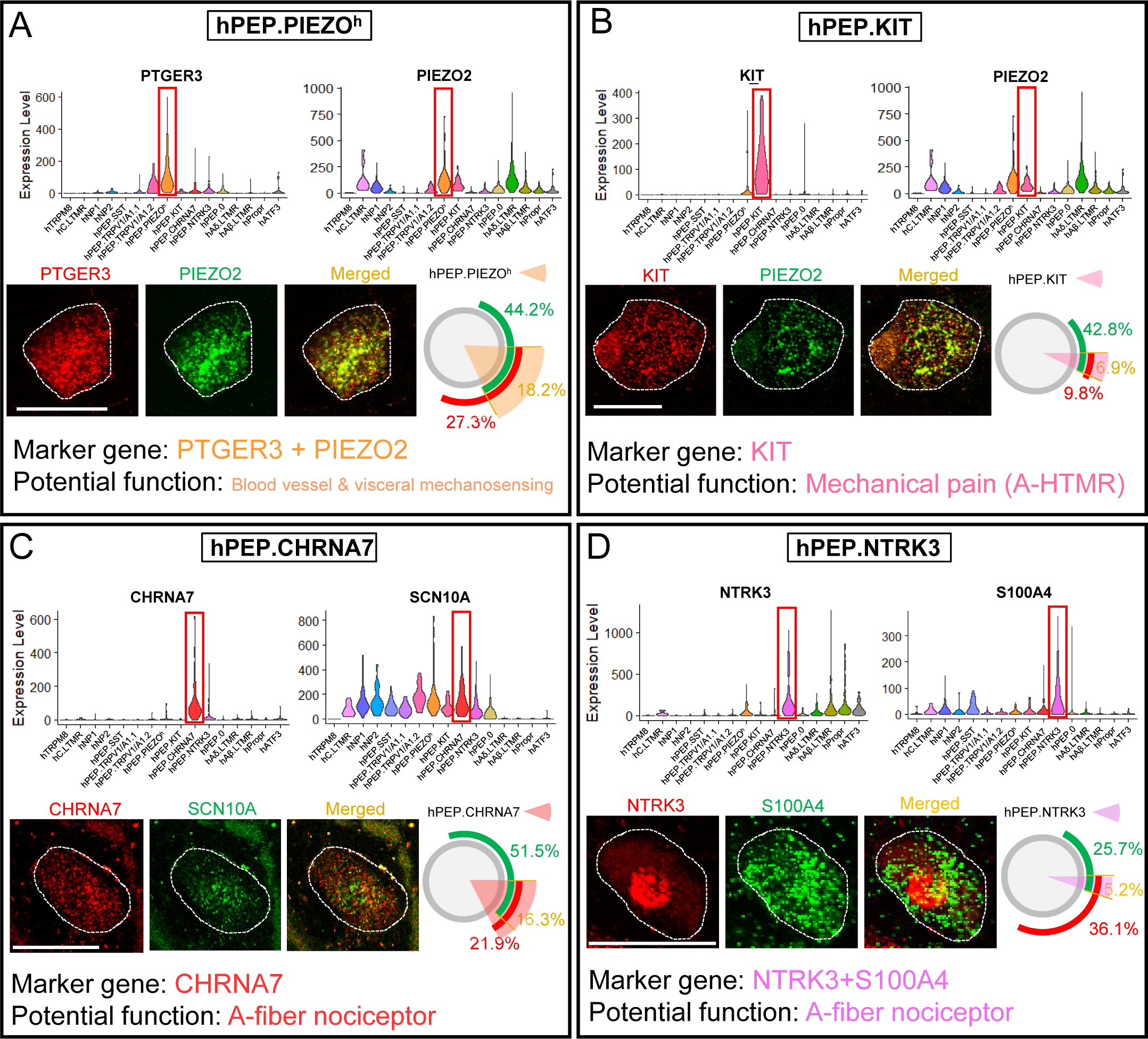
Marker gene expression and validation in A-fiber nociceptors. (**A-D**) Marker genes for specific labeling of each cluster and validation by multiplex FISH for hPEP.PIEZO2^h^ (A), hPEP.KIT (B), hPEP.CHRNA7 (C), and hPEP.NTRK3 (D). The fluorescent images show the detected transcripts in one example hDRG neuron (cell body outlined by the white dashed line). Circle charts next to images show quantification: the arcs indicate the percentage of neurons positive for the given marker gene over all sampled DRG neurons. The sector shaded areas indicate the approximate percentage of each cell type over the total quantified hDRG neurons. N=2, A (165 neurons total), B (173 neurons total), C (196 neurons total), D (191 neurons total). Scale bar, 50 μm.

The hPEP.KIT cluster had the specific expression of a receptor tyrosine kinase, *KIT*, and a medium to low expression level of *PIEZO2* (Fig. 4B). In mouse DRG neurons, *Kit* is more broadly expressed and found in four peptidergic clusters^16^ (Fig. S9B). Cross-species analysis suggest that this cluster mainly corresponded to the mouse PEP3/CGRP-η and macaque PEP3 population (Fig. 2A-C), which are A-fiber high threshold mechanoreceptors (HTMRs), forming circumferential endings around hair follicles and mediating hair pulling pain^65,66^. A more recent study suggests that mouse CGRP-η neurons mediates mechanosensation from the distal colon^67^. Thus, the hPEP.KIT cluster likely functions as a population of fast-conducting mechano-nociceptors.

The third peptidergic A-fiber cluster, hPEP.CHRNA7, featured abundant expression of *CHRNA7* but almost no expression of *PIEZO2* (Fig. 4C). Cross-species analysis revealed that this cluster corresponds to mouse PEP2/CGRP-ζ and macaque PEP2 populations (Fig. 2A-C), which are also marked by the unique expression of *CHRNA7*^16,18^. Interestingly, this cluster also expressed *PVALB* (Fig. 1F), a molecular marker commonly used for proprioceptors in mouse and human. Retrograde tracing from the mouse gastrocnemius muscle labeled *CHRNA7*^+^ DRG neurons^55^, suggesting that hPEP.CHRNA7 may contain muscle-innervating A-fiber nociceptive sensory afferents.

hPEP.NTRK3 is a population of peptidergic A-fibers with high expression of *NTRK3* and *S100A4* (Fig. 4D). Neurons in this cluster expressed a low level of *PIEZO2* (Fig. 4B). Finally, hPEP0 is a population of peptidergic A-fibers that expressed *CALCA* and a moderate level of *PIEZO2* but lack of other specific marker genes. Potential function of hPEP.NTRK3 and hPEP0 are currently unclear. They could be some types of A-fiber mechano-nociceptor^68^.

### Molecular marker expression and validation of C-LTMRs, A-LTMRs and an ATF3 population

hC.LTMR is the putative human C-tactile nerve fibers. All cross-species neuron cluster comparison methods unequivocally identified C-LTMRs as conserved across all three species. Nevertheless, the specific marker gene *CASQ2* for hC.LTM (Fig. 5A) is not detected in either mouse or macaque C-LTMRs^15,18^. Conversely, mouse C.LTMR cells are characterized by exclusive expression of tyrosine hydroxylase (TH) and *SLC17A8* (*VGLUT3*)^15^ (Fig. S9B), which are barely expressed in hDRG neurons (Fig. S9A). Thus, the molecular markers to identify C.LTMRs are different between human and mouse. On the other hand, human, mouse, and macaque C.LTMRs all had conserved expression of GFRA2, another co-receptor of RET, and a zinc finger transcription factor ZNF521 (Fig. S9A-C). Multiplex FISH confirmed that *CASQ2*^+^ cells expressed high levels of *PIEZO2* (Fig. 5A). hC.LTMRs likely mediate innocuous affective touch sensation^70–72^.

**Figure 5.**
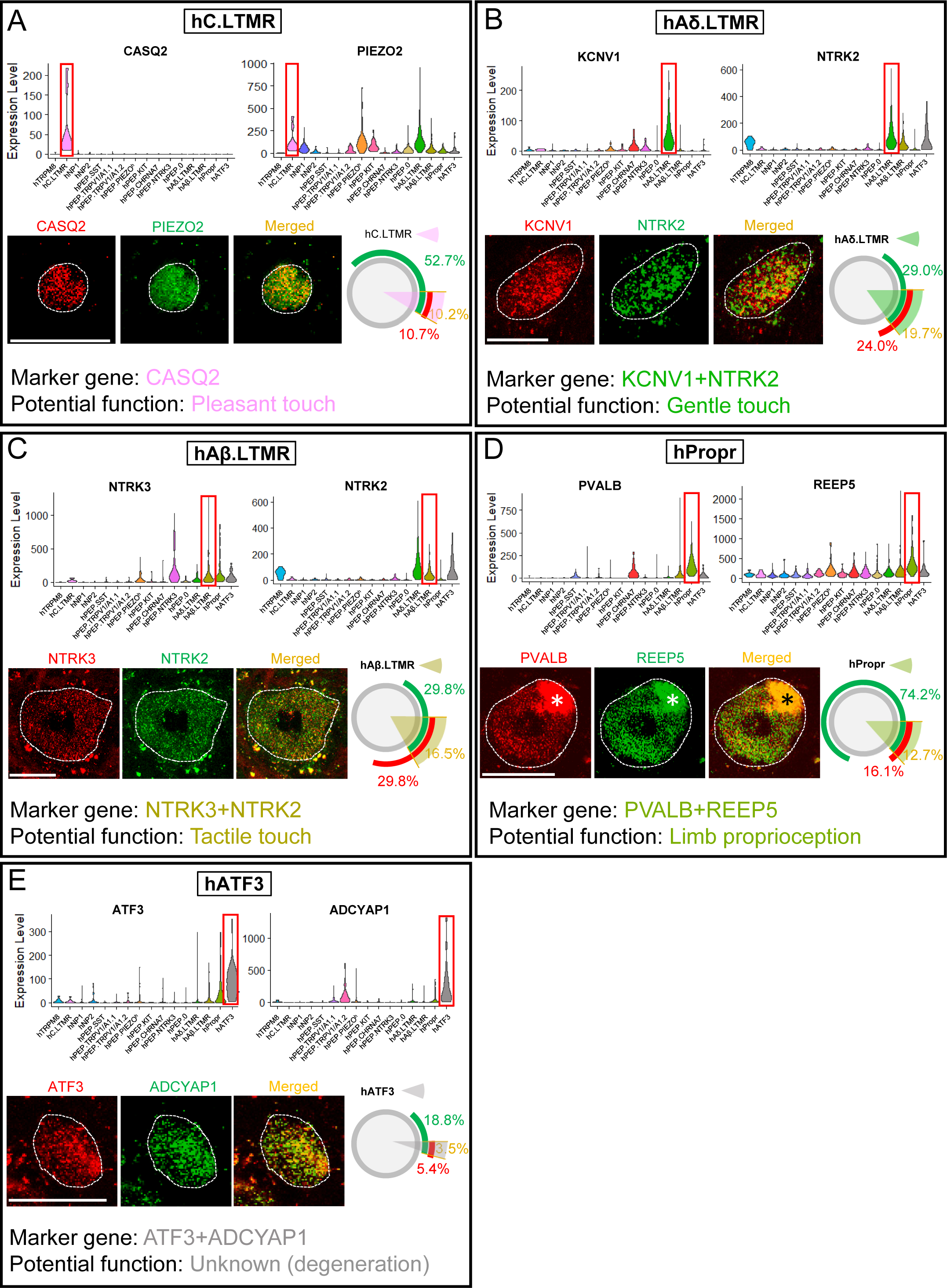
Marker gene expression and validation in C- and A-LTMRs. (**A-D**) Marker genes for specific labeling of each cluster and validation by multiplex FISH for hC.LTMR (A), hAδ.LTMR (B), hAβ.LTMR (C), hPropr (D) and hATF3 (E). The fluorescent images show the detected transcripts in one example hDRG neuron (cell body outlined by the white dashed line). Circle charts next to images show quantification: the arcs indicate the percentage of neurons positive for the given marker gene over all sampled DRG neurons. The sector shaded areas indicate the approximate percentage of each cell type over the total quantified hDRG neurons. N=2, A (205 neurons total), B (183 neurons total), C (188 neurons total), D (198 neurons total), E (202 neurons total). Scale bar, 50 μm.

A-LTMRs were featured by large diameter somas and high expression of *NTRK2* and *NTRK3* (Fig. 5B-C) but a lack of expressions of *SCN10A* and *PRDM12* (Fig. 4C & 1E), two genes highly associated with human nociception^30,73,74^. We identified 4 clusters in this category. hAδ.LTMR was named based on its high expression level of *NTRK2* and *PIEZO2*, but low level of *NTRK3* (Fig. 5A-C), a molecular feature similar to the mouse Aδ-LTMRs (Fig. S9B). *KCNV1* was enriched in this cluster and could serve as a novel molecular marker for identifying this population in hDRG neurons (Fig. 5B). hAβ.LTMR, for tactile touch, expressed higher level of *NTRK3* but a lower level of *NTRK2* compared to hAδ.LTMRs (Fig. 5C). hPropr, for limb position sensing, expressed a high level of a proprioceptor marker *PVALB* and *REEP5* (Fig. 5D).

We also identified a cluster named as hATF3 (Fig. 5E), which contained mainly large diameter neurons and strongly corresponded the “unknown” cluster first identified in normal male mice by the Sharma RNA-seq dataset^16^. This cluster in both human and mouse datasets showed a very conserved molecular profile (Fig. S7A). One of the markers it expressed, *ATF3*, is a transcriptional factor associated with sensory afferent injury, indicating that these cells may represent neurons that are undergoing (or previously underwent) stress/damage/insult (Fig. 5E). *ATF3*^+^ neurons also expressed high level of neuropeptide *ADCYAP1* (Fig. 5E). Since there were no medical records indicating obvious nerve injuries in our human donors, and since the mouse data came from naïve mice, we speculate that these neurons might reflect a very low level of sporadic sensory afferent injury accumulated from daily lives. On the other hand, we cannot exclude the possibility that *ATF3*^+^ cluster represent a population of normal DRG neurons.

### The single-soma deep RNA-seq dataset provides novel insights into the molecular and cellular mechanisms underlying human sensation of itch and pain

Given the high number of unique transcripts per neuronal soma, our dataset is uniquely powerful for molecular discovery. We identified a much higher number of membrane proteins, such as GCPRs, ion channels, chemokine receptors, as well as neuropeptides, compared to 10x Visium Spatial RNA-seq or single-nucleus RNA-seq datasets (Fig. 6A). These discoveries provided novel insights into understanding molecular and cellular bases of physiological recording and psychological experiment results regarding human itch and pain sensation.

**Figure 6.**
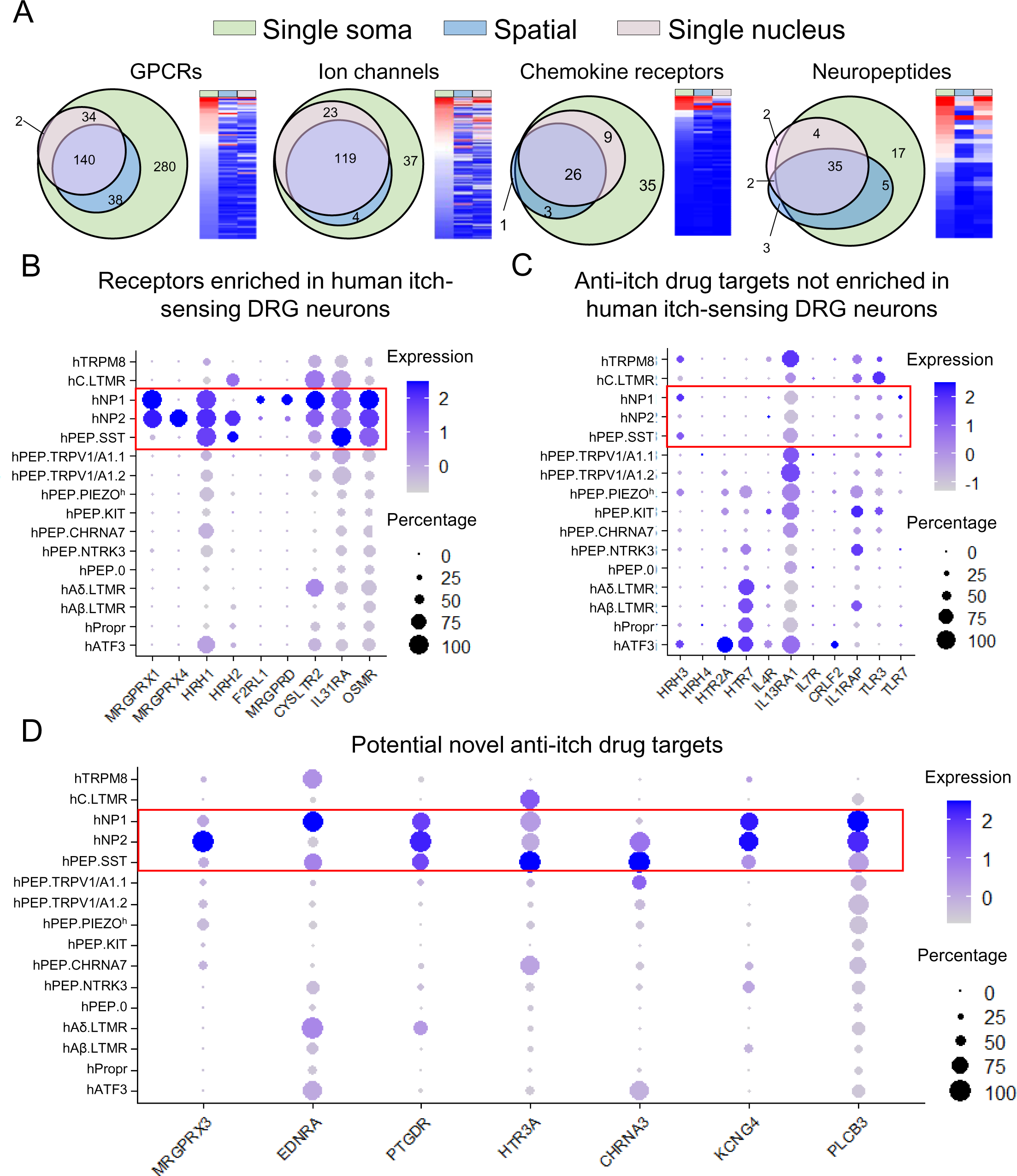
Single-soma deep RNA-seq dataset is powerful for molecular discovery. (**A**) Comparison of the total detected number of GPCRs, ion channels, chemokine receptors, and neuropeptides in single-soma, spatial, and single-nucleus RNA-seq datasets. **(B-C)** Expression of putative itch receptors in hDRG neurons. Some receptors are highly enriched (B), while some receptors are barely detected (C) in human itch populations. (**D**) Novel GPCRs, ion channels, and other genes enriched in human itch populations.

Physiological recordings have identified at least two groups of C-fibers responding to different pruritogens. One group responds to cowhage, a plant triggering intense itch in human^75^, and comprises mechano-sensitive polymodal units^43^. The other group responds to histamine with sustained discharges but is mechanically insensitive^43^. However, the molecular and cellular bases for these observations remain unclear. To understand these physiological properties, we analyzed the expression of histamine and cowhage receptors, and the mechanoreceptor *PIEZO2* in the three potential itch populations, hNP1, hNP2 and hPEP.SST. *PIEZO2* was highly expressed in hNP1, with low expression in hNP2 and almost no detectable expression in hPEP.SST (Fig. 5A). Protease-activated receptor *F2RL1*, the receptor mediating cowhage induced itch in humans^75^, was exclusively expressed in hNP1 (Fig. 6B). Thus, the hNP1 population likely contained C-fiber pruriceptive afferents sensitive to both cowhage and mechanical stimuli. For histamine receptors, *HRH1* was expressed in all three itch populations, *HRH2* was expressed in hNP2 and hPEP.SST but not in hNP1, *HRH3* had low expression in hNP1 and hPEP.SST, while *HRH4* was not detected (Fig. 6B-C). Thus, hNP2 and hPEP.SST clusters are good candidates for histamine-sensitive but mechano-insensitive itch-sensing C-fibers.

When intracutaneously applicated into human skin, some chemicals tend to trigger itch, such as histamine and prostaglandin E2 (PGE2), while others preferentially induce pain, such as serotonin and bradykinin^76^. To explore the potential underlying mechanisms, we examined the expression of all known receptors for these chemicals in our dataset. We found that the receptors of itch-inducing chemicals are enriched in itch sensing populations. For example, histamine receptors *HRH1* and *HRH2*, and PGE2 receptor *PTGER2* were enriched in hNP1, hNP2 and hPEP.SST putative itch-sensing neurons (Fig. S10A1). In contrast, serotonin receptors, *HTR1B* and *HTR1F,* and bradykinin receptor *BDKRB2* were enriched in putative nociceptive populations (Fig. S10A2). The differential expression patterns of these receptors might explain the different sensory experience induced by these chemicals.

### Utility of single-soma deep RNA-seq dataset for discovering novel drug targets and for obtaining insights into molecular targets of existing drugs

Our deep sequencing dataset can also provide novel insights into the mechanisms of existing clinical drugs or for identifying potential novel drug targets. The expression patterns of promising drug targets, such as GPCRs, ion channels, chemokine and cytokine receptors, and neuropeptides, were analyzed (Fig. S11-S14). Here we analyzed itch-related receptors and molecules as an example. A series of itch-sensing receptors have been identified in model organisms^3,46,79^, but which of these targets can be translated remains to be determined. A subset of these itch receptors were indeed enriched in hNP1, hNP2 and hPEP.SST populations, such as the chloroquine receptor *MRGPRX1*, bile acid receptor *MRGPRX4*, histamine receptor *HRH1*, leukotriene receptor *CYSLTR2*, and interleukin receptors *IL31RA* and *OSMR* (Fig. 6B)^16,18^. However, some suspected itch receptors did not exhibit expression enrichment in human itch populations. For example, *HRH4* was reported to mediate histamine-dependent itch in mice^80^, but expression of *HRH4* was barely detected in human itch-sensing populations (Fig. 6C), suggesting that *HRH4* might not be directly involved in human histaminergic itch. The same was found true for *IL7R* and *TLR7*^79^, which were proposed to mediate non-histaminergic itch in mice (Fig. 6C). In addition to these known players, we identified other membrane receptors and signaling molecules in human itch populations (Fig. 6D), including MRGPRX3, EDNRA, PTGDR, HTR3A, CHRNA3, KCNG4, and PLCB3, which could represent novel anti-itch targets.

Gabapentin was originally developed for treating epilepsy and more recently used in the treatment of neuropathic pain and chronic itch^81^. It inhibits neurotransmitter release by acting on α2δ-1 and α2δ-2 voltage-dependent calcium channels *CACNA2D1* and *CACNA2D2*^81^. We found that both receptors were broadly expressed in hDRG neurons (Fig. S10B), suggesting that one potential mechanism by which gabapentin could provide clinical benefit is through inhibiting synaptic transmission of primary afferents, as has been shown in mice^82^.

Opioids and derivatives activate opioid receptors to modulate pain and itch. Agonists of the μ-opioid receptor (*OPRM1*) alleviate pain but elicit itch in humans and model organisms, whereas agonists of the κ-opioid receptor (*OPRK1*) inhibit itch in humans^83,84^. Unlike mouse DRG neurons, which did not5 display cell-type enriched expression patterns of opioid receptors (Fig. S10C), we found that opioid receptors in hDRG neurons were present in some but not other neuron populations. Transcripts of δ-opioid receptor (*OPRD1*) were preferentially expressed in itch populations hNP1 and hNP2, while *OPRM1* was enriched in all hPEP clusters (Fig. S10C). Since opioid receptors are inhibitory GPCRs, our results suggest that activation of *OPRM1* could directly inhibit human nociceptive afferents, while *OPRD1* could be a molecular target for inhibiting itch transmission. On the other hand, *OPRK1* was barely detected in our dataset, suggesting that *OPRK1* agonists may relieve itch through indirect or central mechanisms (Fig. S10C).

### Immunostaining of sensory fibers in the human skin using molecular markers identified by single-soma deep RNA-seq dataset

Although human peripheral sensory afferents can be identified and visualized using the pan neuronal marker antibody against PGP9.5, different types of human sensory afferents cannot be distinguished due to a lack of specific molecular markers. Developing an antibody panel to label different types of human sensory afferents would be invaluable for basic research, translational studies, and diagnostics. To start this direction and to test whether some molecular markers identified by the single-soma RNA-seq are useful for labeling specific types of human sensory fibers, we conducted immunostaining of sensory afferents with leg skin biopsies from three normal adult donors (see Methods for detailed donor information).

*SST* is specifically expressed in hPEP.SST neurons (Fig. 3B). Consistent with this molecular prediction, we found SST^+^ sensory fibers in the human skin sections in the dermis, the epidermis-dermis junction, entering the epidermis (Fig. S15A-C), and near the hair follicle (Fig. S15D-F). In addition, double immunostaining revealed that SST^+^ sensory fibers were CGRP^+^ and made up a subset of CGPR*^+^* sensory fibers (Fig. S15A-F). These results confirmed that human SST^+^ sensory afferents had CGRP proteins and belonged to the PEP but not NP afferents. Since CGRP involves in neuro-immune interactions and other physiological functions^85^, addition of this neuropeptide in human SST^+^ afferents suggests a potential gain of new functions during evolution.

hPEP.KIT neurons specifically express *KIT* transcripts. Strong KIT signals were observed in continuous regenerative cells at the basal layers of the skin, sweat ducts, and hair follicles, serving as a positive control. A few KIT^+^ and PGP9.5^+^ sensory fibers were observed around hair follicles (Fig. S15G-I). Consistent with our RNA-seq results, KIT^+^ sensory fibers were also positive for CGRP and NEFH (Fig. S15J-O), suggesting that they were peptidergic A fibers. In short, these results support that our high-fidelity hDRG neuron single-soma transcriptome dataset is useful for selecting specific molecular markers to label and visualize different types of peripheral somatosensory afferents.

### A novel strategy to study functions of human somatosensory afferents using single-soma deep RNA-seq dataset-informed microneurography recordings

Molecules profiles and physiological properties of hDRG neurons are intrinsically linked: the expressed molecules form the physical basis for physiological properties. Physiological recordings of human somatosensory afferents have been conducted for more than half a century, which generated important parts of our knowledge on the human somatosensory system. Physiological properties of somatosensory afferents generate an important foundation for informing molecular cloning of critical receptors, such as TRPV1 and PIEZO2. Due to a lack of clear molecular compositions of human somatosensory afferents, the reverse experimental direction, from molecular profiles of human sensory afferents to inform physiological recordings, has not been possible. With the single-soma deep RNA-seq dataset, we were finally in a position to do so. Here we focused on two populations of human sensory afferents to demonstrate the feasibility and power of using single-soma deep RNA-seq informed microneurography recordings to study the function of human afferent subtypes.

As mentioned previously, the hPEP.KIT population was highly correlated to the mouse PEP3/CGRP-η cluster^15,18^ (Fig. 2A-C), a population of TRPV1-negative A-fiber fast-conducting hair pull-sensitive mechano-nociceptors^86^. Our single-soma deep RNA-seq data revealed that hPEP.KIT neurons express NEFH and PIEZO2, but not TRPV1 (Fig. 7A), suggesting that they were A-fiber mechano-nociceptors without heat/capsaicin sensitivity. Interestingly, our dataset also revealed that hPEP.KIT neurons expressed a low level of *TRPM8* (Fig. 7A), a feature that has not been reported in mice. Multiplex FISH validated the co-expression of *KIT*, *PIEZO2* and *TRPM8* in hDRG neurons (Fig. S16A). Altogether, these molecular profiles suggest that hPEP.KIT neurons may be A-fiber HTMRs, responsive to cooling but not heating. Thus, we hypothesized that there exist some previously uncharacterized fast-conducting A-fiber HTMRs in the human skin, which can be activated by cooling but not heating stimuli.

**Figure 7.**
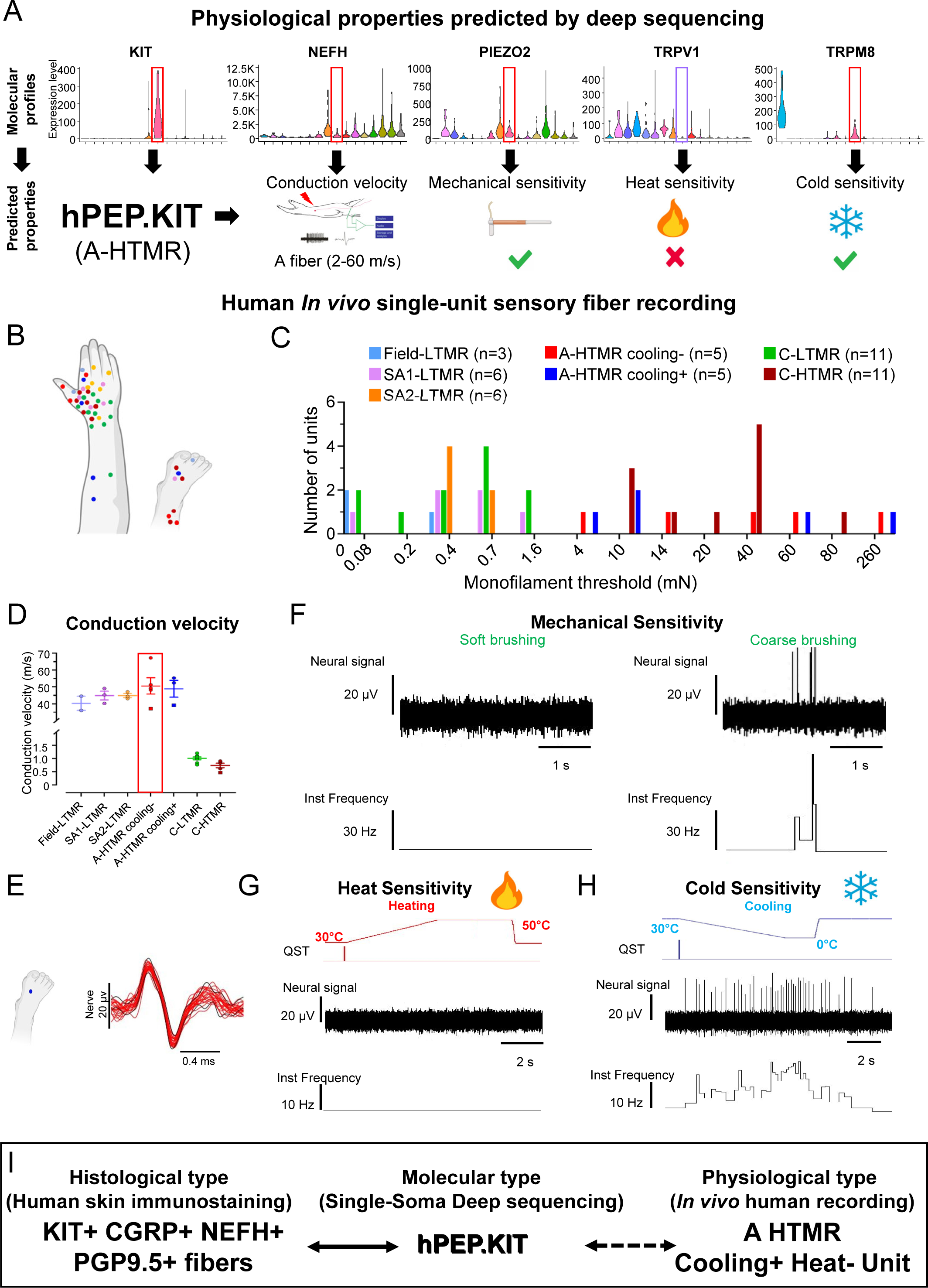
Molecular profile-informed single-unit microneurography recordings revealed novel physiological properties of a population of human A-HTMRs. (**A**) Predicted physiological properties of hPEP.KIT sensory afferents based on single-soma deep RNA-seq data. (**B**) Receptive field locations of recorded single afferents from superficial peroneal (S. peroneal), posterior antebrachial cutaneous (PABCN), and radial nerve recordings (n = 47). (**C**) Distribution of mechanical (monofilament) thresholds for HTMRs and LTMRs in the recorded samples. (**D**) Individual and mean (±SEM) conduction velocities of different HTMR and LTMR types in response to surface electrical stimulation from upper and lower limbs (Field-LTMR: 40.3 ± 4.2 m/s, n=2; SA1-LTMR: 44.9 ± 2.6 m/s, n=3; SA2-LTMR: 44.9 ± 1.2 m/s, n=3; A-HTMR cooling-: 50.6 ± 4.8 m/s, n=5; A-HTMR cooling+: 48.9 ± 5.0 m/s, n=3; C-LTMR: 1.0 ± 0.05 m/s, n=8; C-HTMR: 0.7 ± 0.08 m/s, n=5). (**E**) Spike activities of a putative hPEP.KIT unit (A-HTMR cooling+) in response to repeated stimulations of the receptive field, superimposed on an expanded time scale. (**F-H**) Responses of an A-HTMR cooling+ unit to soft and coarse brushing (F), heating (G) and cooling (H). (**I**) Schematic showing the link between the histologically identified KIT^+^/CGRP^+^/NEFH^+^ sensory afferents and the molecularly defined hPEP.KIT population, likely representing a type of heat/capsaicin-insensitive but cold-sensitive A-HTMRs.

To test this idea, using the *in vivo* electrophysiological technique of microneurography, single-unit axonal recordings were performed from the radial, antebrachial, and peroneal nerves of healthy participants (Fig. 7B-D). The A-HTMRs (n=10) were identified by their insensitivity to soft-brush stroking while responding to a rough brush and displaying high indentation thresholds (≥4 mN); further, they had Aβ-range conduction velocities (>30 m/s, Fig. 7B-F). Remarkably, a subtype of these heat-insensitive A-HTMRs (n=5) units responded to cooling (Fig. 7G-H). Compared to mechanically evoked responses, those evoked by cooling although relatively modest but reproducible with tested in triplicates for each recording (Fig. S16B). Furthermore, these cooling-evoked responses persisted during the sustained phase (Fig. S16B). These observations confirmed our prediction that some human cutaneous A-HTMRs are cold-sensitive but heat-insensitive. Based on the transcriptome dataset and current knowledge about molecular receptors for heat and cold, hPEP.KIT seems to be the only population that has the molecular basis for this distinct combination of physiological properties. Thus, we propose that the molecularly defined hPEP.KIT population is correlated to the physiologically defined A-HTMRs with cold but not heat sensitivity (Fig. 7I).

Human C-LTMRs are readily found during microneurography recordings from the upper limb including the distal regions^87^, but they seem to sparsely innervate the distal lower limb^88^. This is consistent with our sequencing results wherein the hC.LTMRs constituted a small population of neurons in lower thoracic and lumbar level DRGs (Fig. 1G). Unexpectedly, our sequencing data revealed that human hC.LTMRs had almost no expression of the cold and menthol receptor *TRPM8* (Fig. 8A) though C.LTMRs display sensitivity to cooling in both humans and mice^89,90^. Another unexpected finding was that hC.LTMRs expressed *TRPV1* (Fig. 8A). Multiplex FISH confirmed that *CASQ2*^+^ hC.LTMRs were TRPM8^−^ but TRPV1^+^ (Fig. S17A). This expression pattern suggested that hC.LTMRs might respond to heating and capsaicin, a novel physiological property that has not been discovered in rodent or non-human primate models, but is predicted not to respond to menthol. In human microneurography, the C-LTMRs (n=11) were identified by their soft-brush sensitivity, low indentation threshold (in this case, ≤0.7 mN), and slow conduction velocity (∼1 m/s, n=11, Fig. 7B-C, 8B-D). Consistent with mouse C-LTMRs, these human counterparts responded to hair movement (Fig. 8D) and dynamic cooling (Fig. 8E). In two of them, after having confirmed the cooling response, we applied menthol to their individual receptive fields resulting in a cold sensation, but the recorded C-LTMRs, while still responsive to mechanical and thermal stimuli, were not activated by menthol (Fig. 8F). Remarkably, microneurography recordings showed that a subset of human C-LTMRs responded to dynamic heating (5 out of 11 units or 45%, example shown in Fig. 8G). This proportion of heat-sensitive C-LTMRs aligns with findings in the rabbit (8 out of 18 units or 44%)^91^. In three C-LTMRs, after having confirmed the heating response, we applied capsaicin to their receptive fields. Consistent with the sequencing results, all three were activated by capsaicin (Fig. 8H). Relative to mechanical stimulation, the C-LTMR responses to dynamic temperature changes were comparatively modest. However, these responses were reproducible, having been tested in triplicates for each modality and recording (Fig. S16B-C). Interestingly, the C-LTMR has different cooling response properties compared to A-HTMR cooling+ fibers. The cooling-evoked responses rapidly diminished in C-TLMR but persisted during the sustained phase in A-HTMR cooling+ fibers. The existence of a polymodal (mechano-heat-cold) C-LTMR type is novel and confirms the sequencing predictions. Furthermore, the cooling response must be mediated through a non-TRPM8 mechanism.

**Figure 8.**
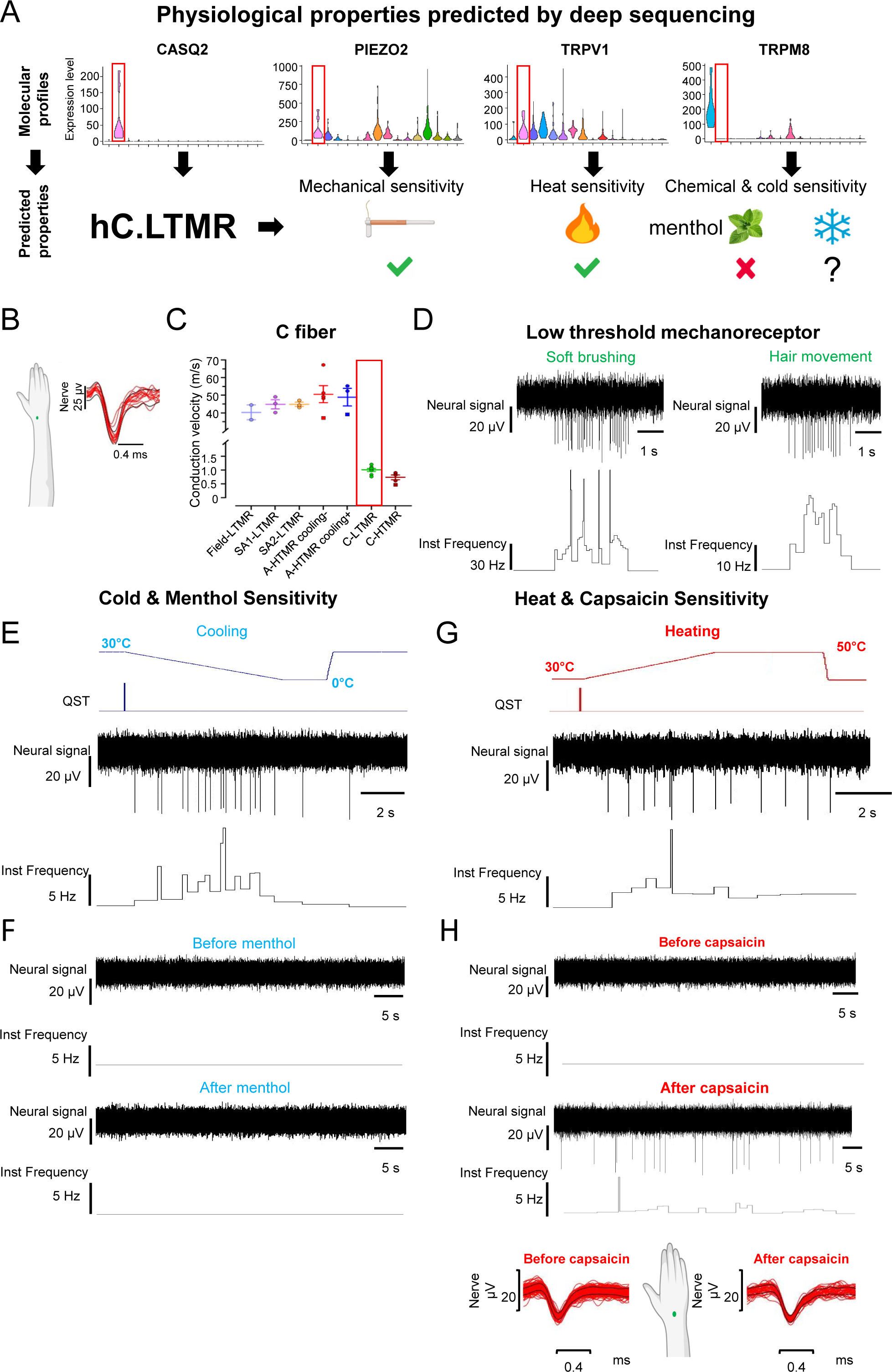
Molecular profile-informed single-unit microneurography recordings revealed novel physiological properties of a population of human C-LTMRs. (**A**) Novel physiological properties of hC.LTMR sensory fibers predicted based on gene expression obtained from the single-soma deep sequencing. (**B**) Spike activity of a hC.LTMR unit in response to repeated stimulations of the receptive field, superimposed on an expanded time scale. (**C**) Individual and mean (±SEM) conduction velocities of different HTMR and LTMR types in response to surface electrical stimulation from upper and lower limbs (the same plot from Fig. 7D) (**D-H**) Responses of a hC.LTMR unit to soft brushing and hair movement (D), cooling (E), menthol (F), heating (G) and capsaicin (H). Spike activity of that hC.LTMR before and after capsaicin application, overlaid on an expanded timescale (H). Conduction delay was adjusted based on the latency of electrically triggered spiking for that recorded afferent. Note, different scaling.

For comparison, our recordings also identified C-HTMRs (n=11) that did not respond to soft-brush stroking and hair movement (they responded to a rough brush), had high indentation thresholds (in this case, ≥10 mN), and slow conduction velocities (∼1 m/s, Fig. 7B-D & S17B-E). Based on their temperature responses (tested in 9 units), a mix of C-fiber mechano-heat (n=6), C-fiber mechano-cold (n=2), and C-fiber polymodal (mechano-heat-cold, n=1) subtypes were identified. An example of heating and capsaicin responses of a C-fiber mechano-heat nociceptor is shown in Fig. S17F-G. Collectively, these results highlight the accuracy and utility of our single-soma deep RNA-seq dataset of hDRG neurons in guiding and informing the functional characterization of human somatosensory afferents and the power of combining the two approaches. We believe that this signifies a promising new research direction to link molecular and physiological types and to discover novel functional properties of human somatosensory afferents.

## Discussion

Despite the high relevance to human health, molecular and cellular mechanisms underlying normal and pathological human somatosensation remain largely elusive. In this study, we developed a novel LCM-based approach for single-soma deep RNA-seq and recovered more than 9000 unique transcripts per neuron over 1000 adult hDRG neurons. The sequenced hDRG neurons were clustered into 16 molecular groups, displaying similarities to and remarkable differences from macaque and mouse DRG neurons. As exemplified in the study, our dataset provides important and novel insights into human pain and itch sensory phenomena, explains mechanisms of drug effects, and represents a rich resource to identify new molecular targets for modulating the activity of itch- and pain-sensing primary afferents. Moreover, our dataset successfully guided histological labeling (Fig. S15) and physiological recording (Fig. 7, 8 & S17) of different types of human sensory afferents, leading to discovery of their new anatomical features and physiological properties, and serving as a common ground to connect them all together (Fig. 7I).

Single-cell RNA-seq of human DRG/TG neurons has been technically challenging. One main hurdle is to isolate sensory neurons from a large population of non-neuronal cells. The traditional enzymatic and mechanical dissociation method is incompatible with human DRG/TG neurons. Strategies, including spatial transcriptomics and single-nucleus RNA-seq^22,23,92^, have generated some pioneering datasets characterizing the molecular profiles of human DRG and TG neurons. Nevertheless, the number of transcripts, sequencing depth, or single-cell resolution of the previous studies needs to be improved. Importantly, though nuclear and soma transcripts are intrinsically linked and overlapping to some extent, the soma contains much more transcripts both quantitatively and qualitatively. Soma transcripts are also one step closer than nuclear transcripts for functions. Thus, soma transcripts are preferable, if available, for cell type clustering and functional interpretation. In this study, we developed a new strategy by combining fresh frozen hDRG tissues, cryo-section, laser capture microdissection of individual neuronal soma, and Smart-seq2 deep sequencing. Fresh frozen tissue and cryosection techniques minimized transcriptomic changes during the tissue transportation and single-cell isolation process. LCM allowed isolation of DRG neuronal soma with minimal contamination from surrounding nonneuronal cells, while preserving information about cellular morphology, and the Smart-seq2 protocol enabled a high recovery rate of mRNA molecules. Although LCM has been used for isolating a group of neurons for RNA-seq^6,7^ or single TG neurons for genomic DNA analysis^93^, the successful application of LCM for single-cell transcriptomic analysis has not been achieved before. Thus, we have established a new method for single-cell RNA-seq of adult hDRG neurons. Our approach should be readily applicable to other human neurons with large soma sizes, such as other peripheral ganglia neurons, motor neurons, etc.

Different single-cell RNA-seq approaches, including single-nucleus RNA-seq^21,23,92^, spatial transcriptomics^22^, and our LCM-based single-soma RNA-seq, have generated four datasets of transcriptome profiles and cell type clusters of hDRG neurons. These datasets and results overlap to some extent but also exhibit some significant differences (an example shown in Fig. S4). The observed differences are likely caused by both biological and technical features associated with each method. Given the high sequencing depth of transcripts from the neuronal soma, our approach is powerful for molecular discovery, especially for functional molecules expressed at a low level. For example, our approach detected the specific expression of *MRGPRX1 and MRGPRX4*, two important pruritogen-sensing GPCRs, in hNP1 and/or hNP2 neurons, while the previous datasets barely or not detected these transcripts^22,23^. Our analysis of the expression of receptors, ion channels, and neuropeptides, in human itch-sensing DRG neurons have also identified a set of potential new targets for modulating activities of these sensory afferents (Fig. 6).

In addition, our sequencing and microneurography results raise many interesting questions regarding the molecular receptors and afferent types involved in human cold and mechanical pain sensation. For example, human hC.LTMRs rely on a non-TRPM8 cold receptor or another mechanism for their cooling sensitivity^88^. We noticed a low expression level of *TRPA1*, which also has cold sensitivity^94,95^, within this population (Fig. S7E). Whether *TRPA1* or some currently unknown cold receptors mediate cooling sensitivity in this population will be of interest for future research. The discovery of the hPEP.KIT population indicated a potential role for PIEZO2 in human mechano-nociception. This population also responded to cooling, which is a curious property of a large-diameter myelinated nociceptor. Our discovery of TRPM8 expression in these neurons provides a molecular explanation for this unique property. In addition, a likely molecular type for C-HTMRs, which display a mixture of responsiveness to mechanical forces and temperature, is the hPEP.TRPV1/A1.1 population. Since hPEP.TRPV1/A1.1 neurons have no expression of *PIEZO2* or *PIEZO1* (Fig. S9A), a non-*PIEZO* mechanoreceptor may exist in these neurons for mediating mechanical pain sensation. This is consistent with reports that patients with *PIEZO2* loss-of-function mutations still have normal mechanical pain threshold and sensitivity^68,96^. One suggested mechanical pain channel, TMEM120A (TACAN), is broadly expressed in all types of hDRG neurons (Fig. S9A). This expression pattern does not support its purported role as a mechanical pain receptor in humans. Both C-LTMRs and A-HTMRs responded to cooling in human microneurography. While the responses were comparatively modest in contrast to mechanically evoked responses, they were consistent on a trial-to-trial basis. This suggests that the requisite circuity may already be in place, which could potentially have implications for understanding thermal hypersensitivities in pathological states. Indeed, there is indirect evidence from human psychophysics and targeted pharmacology, pointing to the role of C-LTMRs in mediating acute cold allodynia^97^ and the role of A fibers in signaling chronic cold allodynia^98^. In short, our dataset serves as an important atlas for understanding sensory properties of hDRG afferents and somatosensation more generally.

Insights into the relationship of the neuronal types between the mouse and human are essential, as such knowledge is important for translation of pre-clinical discoveries and for inferring functions of human sensory afferents. The three different analysis strategies used in this study complement each other as they have different strength and weakness (see Methods for details). To our satisfaction, despite the different advantages and disadvantages, results from these methods were highly consistent with each other (Fig. 2 & S5). Our results suggested that many broader functional groups of DRG sensory afferents are conserved across species, despite noticeably molecular differences (Fig. 2 & S6). The greatest divergence between mouse and hDRG neurons was observed among C- and A-fiber nociceptors. Mice contain four C-fiber nociceptors (PEP1 neuron subtypes), two expressing TRPV1 but not TRPA1 or PIEZO2 (PEP1.1 and PEP1.2) and another two which expresses all three channels (PEP1.3 and PEP1.4) (Fig. 2A & 2C). While not much is known about mouse PEP1.2/CGRP beta, PEP1.1 represents a noxious heat sensory type, terminating with free nerve endings in epidermis of hairy skin^99^. PEP1.3/CGRP gamma neurons innervate mainly internal organs and are a silent nociceptor type, becoming active during inflammation^100^. No functional studies have been performed on PEP1.4 in the mouse. These neurons are speculated to represent C-HTMRs which are functionally known to exist in the mouse^101^. In humans, it seems that this diversity of C-fiber nociceptors has conflated into two types, the hPEP.TRPV1/A1.1 and hPEP.TRPV1/A1.2 (Fig. 2A & 2C), expressing both TRPV1/TRPA1 with or without PIEZO2, respectively. If similar to mice, different neuronal populations in human are somewhat specialized according localization of peripheral target, one of these types may represent deeply innervating silent nociceptors, known to be present in human^102^, and the other involved in slow noxious thermal and burning cutaneous sensation and perhaps joint pain. One thing to note is that our dataset contains transcriptomes of hDRG neurons from the lower thoracic and lumbar levels, which mainly convey sensations from the leg and lower back. Thus, it remains an open question whether humans have a population of internal organ-specific C-nociceptors, like the mouse PEP1.3 cluster. This question will be resolved when sequencing more hDRG neurons from the thoracic level.

For A-fiber nociceptors, only two types have been identified in mice. The PEP2/CGRP-zeta mechano-heat nociceptive population expresses PIEZO2 and low levels of TRPV1^16,33^, consistent with heat activation only at very high temperatures^99^, and the PEP3/CGRP-eta mechano-nociceptive conveying fast and sharp pain, including hair pulling^44,103^. In this study, we identified five types A-fiber nociceptors in hDRG neurons (Fig. 1G, 2A & 2C). Two populations have clearly conserved features to mouse: hPEP.CHRNA7 showed molecular similarities to mouse and macaque PEP2/CGRP-ζ, and hPEP.KIT to mouse and macaque PEP3/CGRP-η. The other three populations displayed much greater divergence (i.e. hPEP.NTRK3, hPEP.PIEZO^h^, and hPEP.0). The hPEP.PIEZO^h^ cluster is particularly interesting since these neurons express very high levels of *PIEZO2* and *PIEZO1,* unlike any neuron types in the mouse. Our results suggest that fast pain and interoception might have been an evolutionary preference in humans as compared to rodents, an effect that may relate to significantly increased human body sizes.

Finally, in this study, we sequenced over 1000 DRG neurons from 6 DRGs at the thoracic (T11-T12) and lumbar (L2-L5) levels of three Caucasian human donors with no obvious somatosensory or systematic diseases and substance uses (Supplementary tables 1 & 2). It is about ∼150 to 200 neurons per DRG, which represent 1-2% of the total neurons within a hDRG. Thus, sequencing more neurons might give an improved representation. The existence of different types of RA and SA Aβ-LTMRs and different types of proprioceptors in humans is well known, but our current dataset did not have the resolution yet to separate them (Fig. 1G). With increased number of neurons sequenced, we anticipate discovering an even greater heterogeneity among hDRG neurons. In addition, DRGs at different spinal levels innervate different peripheral target tissues^29^. Sequencing DRG neurons from different spinal levels will help to uncover molecular and cellular mechanisms underlying physiological functions. Moreover, increased sampling from additional donors representing different demographics would also be critical for investigating sex, race, and age-related differences. Last but not the least, pathological conditions greatly alter the transcriptomic landscape^104,105^. Systematic comparison of molecular and cellular changes between donors at the baseline condition (like the screening criteria of this study) and those with chronic itch or chronic pain would be of great value, if samples are available, for understanding pathological mechanisms and for identifying molecular targets for effective treatments.

## Acknowledgements

We thank all tissue donors and their families for the generous donations. The new knowledge pieces generated from this study would not be possible without their precious gifts. We thank NDRI for helping with hDRG procurement, Dr. Isaac Chen, Ms. Arianna Unger, and other members at the UPenn (University of Pennsylvania) department of neurosurgery for helping with patient DRG procurement for method optimization, the UPenn Skin Biology and Disease Resource-based Center (SBDRC) for helping with laser capture microdissection, and the Children’s Hospital of Philadelphia (CHOP) Center for Applied Genomics (CAG) for helping with sequencing. We thank Dr. Yulong Li for his encouragement in starting this new research direction and all lab members in the Luo and Ma labs for their helps in conducting this project. This project is supported by an internal funding from UPenn to Dr. Luo. Dr. Luo is also supported by NIH fundings (R01 (R01NS083702), R34 (NS118411), U01 (EY034681), and P30 (AR069589)). Dr. Ernfors is supported by the Swedish Research Council (2019-00761), Knut and Alice Wallenbergs Foundation, ERC advanced grant (PainCells 740491). Dr. Olausson is supported by Knut and Alice Wallenberg Foundation (grants KAW 2019.0047 and KAW 2019.0487). Dr. Nagi is supported by the Swedish Research Council (2021-03054), Swedish Medical Society (SLS), and ALF Grants, Region Östergötland.

## Materials and Methods

### Human tissues and subjects

Human DRG tissues were procured from National Disease Research Interchange (NDRI). The research application was approved by the NDRI Feasibility Committee (RLUW1 01). Six DRGs between T11 to L5 of three human donors aged from 23-61 years were used in this study. The dissected DRG tissues from human donors were immediately imbedded in OCT, shipped to the Luo lab on dry ice and stored in −80 L until use. The information of DRGs and dis-identified donors and screening criteria are summarized in the Supplementary Tables 1 & 2. As determined by the University of Pennsylvania IRB, this study was exempted from the human subject requirements for the Luo lab.

The human skin biopsies were extracted from three healthy donors at the college of medicine, University of Florida. This tissue procurement was approved by the university IRB (protocols IRB201500232 and IRB202300291). The information of human skin biopsies and dis-identified donors is summarized in the Supplementary Table 3. These three donors are members of one family and have no noticeable abnormal somatosensation or peripheral neuropathy. All participants were provided written informed consent and signed the document.

*In vivo* recordings of peripheral sensory afferents of healthy human subjects were performed at Linköping University, Sweden. These subjects were recruited through social media. All participants were provided written informed consent before the start of the experiment. The study was approved by the ethics committee of Linköping University (dnr 2020-04426) and complied with the revised Declaration of Helsinki.

### Laser capture microdissection of hDRG neurons

The hDRGs imbedded in OCT were cryosectioned (Cryostat Leica Cm1950) into 20 μm sections and mounted onto Arcturus PEN Membrane Frame Slides (Applied biosystems, LCM0521). One of every five consecutive sections was collected for laser capture microdissection to avoid repeated dissection of the soma from the same neuron in different sections. The slides were stored in −80 L until further use.

On the day of laser capture microdissection, the slides were transferred to the SBDRC laser capture microdissection (LCM) core on dry ice. Before dissection, the section was briefly stained with RNase free Arcturus™ Histogene™ staining solution (Applied biosystems, 12241-05) for better visualization of neuronal soma: 70% cold EtOH for 30s; Histogene™ staining for 10s; 70% cold EtOH for 30s; 95% cold EtOH for 30s; 100% cold EtOH for 30s and air dry for 2min. Then, the slide was put onto laser capture microdissection microscope system (Leica LMD6) for the neuronal soma dissection. The laser was calibrated, and the laser intensity was adjusted to achieve best dissection efficiency. The dissected individual neuronal soma was collected in the cap of a 200 μl PCR tube containing 4 μl lysis buffer^26^. The sequencing library was generated following Smart-seq2 workflow^26^. The libraries passing through all quality controls were selected for the final sequencing.

### Sequencing and sequence alignment

The libraries were pooled together (384 samples for one batch) and sequenced on NovaSeq 6000 platform by the Children’s Hospital of Philadelphia (CHOP) Center for Applied Genomics (CAG). Raw sequencing data was demultiplexed with bcl2fastq2 v.2.20 (Illumina) followed by Tn5 transposon adapter sequences trimming with Cutadapt^106^. The processed reads were then aligned to human genome (GRCh38 GENCODE as the reference genome, and GRCh38.104. GTF as the annotation) using STAR v.2.7.9a49^107^. STAR quantMode GeneCounts was used to quantify unique mapped reads to gene counts.

### Analysis of single-soma RNA-seq data of hDRG neurons using Seurat and R

R (version 4.1.2) and Seurat (version 4.0.5) were used for the single-cell RNA-seq analysis. Six objects were created from the individual biological replicates. The data were normalized (NormalizeData) after which 4500 most variable features were selected (FindVariableFeatures). To mitigate batch effects between replicates, we used Seurat’s integrated analysis approach that transforms datasets into a shared space using pairwise correspondences (or “anchors”)^19^. Anchors were first defined using FindIntegrationAnchors (dimsL=L1:30) and the data were then integrated (IntegrateData) and scaled (ScaleData), followed by principal component analysis (PCA) (RunPCA, npcsL=L50). For clustering, the final parameters were: RunUMAP, reductionL=Lpca, dimsL=L1:25; FindNeighbors, reductionL=Lpca, dimsL=L1:25; FindClusters, resolutionL=L3.4. Highly similar clusters without clearly distinguishable markers were merged to produce the final 16 clusters.

### Analysis of single-soma RNA-seq data of hDRG neurons using Conos

For Conos^31^ analysis, single-soma hDRG data were integrated using CCA space $buildGraph(kL=L8, k.self=3, spaceL=L“CCA”, ncomps=30, n.odgenes=2000, verbose=TRUE, snn=T, snn.kL=L10). For human single-soma and human single-nucleus dataset, co-integration was performed as $buildGraph(kL=L8, k.self=3, spaceL=L“CCA”, n.odgenes=2000, verbose=TRUE, snn=T, snn.kL=L10). For Conos co-clustering mouse (Sharms) dataset was downsampled to max 300 cells per cluster, co-integration was performed as $buildGraph(kL=L8, k.self=3, spaceL=L“CCA”, n.odgenes=2000, verbose=TRUE, snn=T,snn.kL=L10). Macaque (Kupari, SmartSeq2 dataset) was used for interspecies analysis. For Conos co-clustering macaque/human graph was built as $buildGraph(kL=L4, k.self=3, spaceL=L“CCA”, ncomps = 30, n.odgenes=2000, snn=F, snn.kL=L10). For all UMAP plots in Conos graphs were embeded as: $embedGraph(methodL=L“UMAP”, spread=15, seed = 3). Label propagation ($propagateLabels) was run using method “diffusion”.

### Methods used to elucidate cross-species cluster relationships

We used four different interspecies analysis approaches. First, Conos, which uses graph-based dataset integration and was developed to co-cluster and compare datasets originating from different RNA-seq platforms and species (Fig. 2A-B, S5A-B). Second, probabilistic neural network analysis which is a variant of machine learning in which learning module is trained with one dataset and then testing other datasets for pattern recognition and probability output^18^ (Fig. S5C-D). Third, neural network based hierarchical clustering analysis (Fig. S5E). In the hierarchical clustering analysis, each query neuron types, either human or macaque, is assigned weights of the sensory-type associated patterns by a neural network, which was trained with gene patterns including both species specific and shared cross-species features in the different sensory neuron types. The weighted gene patterns were then used for dimensional reduction and nearest neighbor analysis to infer the hierarchical relationship. Finally, we performed transcription factor associated gene regulatory network analysis across all three species using genes network modules presumably driven by individual transcription factors (Fig.2D). These three methods have different strengths and weaknesses. Conos finds shared principal components between integrated datasets, but some species-specific features may be lost and lead to impaired statistical sufficiency during integration and furthermore can be affected by the number of principal components and nearest neighbor distance. Machine learning is based on gene expression and is not supervised by shared latent space (i.e. common principal components). Each single reference dataset is used to train the machine learning module and then tested by the other dataset. Thus, an advantage of this method is that at the stage of machine learning, datasets are not integrated, and hence, probability calculations are not affected by principal components or nearest distance. Conversely, its disadvantage is that since it emphasizes the features of each cell type, the learning accuracy and reliability depend on the robustness of the reference or training dataset. For the third approach, machine learning-based hierarchical clustering, we extracted the weight of cell type-specific features to construct the latent space covering all cell types across species, whether shared or not. With this strategy, we tried to obtain sufficient latent space, as compared to Conos, by training and predicting every dataset independently and furthermore, parameterization was used to find the most robust hierarchical clustering.

### Cell type probabilistic similarity estimation across cell types and the data integration across species

The assessment of cell type purity, the probabilistic similarity, and cell-type integration across species are performed using packages in a machine learning based single-cell analysis toolkitscCAMEL, released separately at https://sccamel.readthedocs.io/.

### Probabilistic similarity estimation across cell types

The calculation of cell-type probabilistic score has been described in SWAPLINE package^108^. Briefly, a vanilla neural network model was built for cell-type classification. To train the model, we removed the cell cycle–related genes and then computed the most variable features. In addition, we ranked the marker genes for each cell type by two heuristics for the cell-type specificity of both fold change and enrichment score change. Subsequently, the ranked marker genes and the most variable genes were merged, log-transformed, and scaled by min-max normalization for learning models. The frame of the neural network model and the parameters have been described in the SWAPLINE package. The learning accuracy of the neural-network classifier was inspected against epoch numbers and was estimated by k-fold cross-validation (k = 3). The learning rate and learning epochs were selected according to the maximum point of the learning curve reaching the accuracy plateaus. The probabilistic scores from mouse and macaque species against human reference were visualized in violin plot.

### Data integration across species

For the integration task, we applied interpretable neural-network learning. First, we took one dataset from the dataset pool. We trained a neural-network classifier by learning the transcriptional features of each cell-types in this dataset and then calculated the trained cells’ probabilistic scores against all cell-types. Subsequently, we used all other datasets as query datasets and calculated the probabilistic score of every cell in each query dataset via the trained classifier. Then, we took another dataset from the dataset pool and repeated the training and prediction. We repeat the training and prediction till every dataset has been used as a training reference for the predictions. Here, we consider that the probabilistic score of each cell reflects the weighted gene patterns representing each trained cell-type. Thus, we merged the probabilistic scores of all cells from all trained and predicted datasets for the principal component analysis. The most significant principal components were determined by the elbow method and subsequently used as the latent space for further downstream analysis. The tree plot was constructed with the parameter of 11 principal components, 90 nearest neighbors, and correlation metric. The trained cell-type similarity was calculated with the correlation distance and the average/UPGMA linkage and visualized in the hierarchical heatmap.

In parallel, we normalized the gene expression by interpretable learning. We transformed the gene symbols of each species into the nomenclature in Homo sapiens. We estimated the features’ weights in each reference cell-type by using the DeepLift algorithm^109^. The gene expression of each cell that has been learned or predicted in one trained reference dataset, was inferred by the matrix multiplication between the features’ weights and the cell-type probabilistic scores. And the final gene*cell expression matrix was calculated by the average of non-empty values across all datasets. Using this normalized expression matrix, we enriched the mostly co-expressed genes via spearman correlation. These co-expressed genes were used for inferring the TF associated gene patterns via a modified GENIE3, as described in^110,111^. The result was visualized as a hierarchical heatmap.

### Multiplex FISH, confocal microscopy imaging, and quantification

OCT embedded freshly dissected human lumbar or thoracic DRG tissues were cryosectioned at 20Lµm thickness and mounted on glass slides. The slides were stored in −80L°C to preserve RNA integrity until use. RNAscope Fluorescent Multiplex Reagent Kit and RNASCOPE probes for the targeted genes (Advanced Cell Diagnostics Inc.) were used for *multiplex FISH*. RNAscope *in situ* hybridization was performed in accordance with the manufacturer’s instructions. In brief, fresh frozen hDRG sections were fixed, dehydrated, and treated with protease. The sections were then hybridized with the respective target probe for 2 hr at 40°C, followed by two to three rounds of signal amplification. The sections were then mounted under coverslips, sealed with nail polish, and stored in the dark at 4°C until imaged. A Leica SP5 confocal microscope was used to capture images and ImageJ was used for image analysis. In some DRG neurons, accumulation of lipofuscin in part of cells caused strong autofluorescence in all channels. These signals were considered as non-specific background (labeled by asterisk) were excluded for analysis. (See Fig. S9 for examples). The percentage of each cluster over all DRG neurons could be a little bit overestimated due to the following two reasons: 1) Some marker genes or marker gene combination may also label a small subset of other cell types; 2) An underestimation in quantification of total neuronal numbers because some cells have neither multiple FISH signals nor DAPI (4′,6-diamidino-2-phenylindole) nucleus staining signals.

### Human skin biopsy extraction, processing, and immunostaining

Dermal skin punch biopsies were performed as described herein. Briefly, 1 cc of lidocaine was injected subdermally at each biopsy location (Supplementary table 3). A total of six, 3 mm dermal skin punch biopsies were performed on each patient. Excised skin was immediately placed in 1.5 mL eppendorf tubes containing 4 degree Celsius 4% paraformaldehyde (PFA) solution that was freshly prepared on the same day of the skin biopsy procedure. Biopsy tissue was fixed in 4% PFA (dissolved in PBS) for exactly four hours at 4 degrees, followed by 2 X 30 minutes washes in phosphate buffered saline (PBS) solution, and then cryoprotected using 1XPBS, 30% sucrose at 4 degrees. These tissues in cold 1XPBS, 30% sucrose were overnight shipped to the lab of Integrated Tissue Dynamics LLC.

The skin biopsies were mounted in OCT and cryosectioned into 14 μm sections. Adjacent sections were collected by continuous slides. Immunofluorescence in this study was performed using combinations of mouse monoclional anti-human PGP9.5 (Protein Gene Product, CedarLane, Burlington, Canada, 31A3, [source UltraClone Ltd, Isle of Wight, UK]; 1:200), sheep polyclonal anti-human CGRP, mouse monoclonal anti-human NEFH (Sigma [ab142]; 1:400), rabbit antihuman SST (ImmunoStar 20067, Hudson, WI, USA), and anti-human KIT. Slides were preincubated in 1% bovine serum albumin and 0.3% Triton X-100 in PBS (PBS-TB) for 30 minutes and then incubated with primary antibodies diluted in PBS-TB overnight in a humid atmosphere at 4°C. Slides were then rinsed in excess PBS for 30 minutes and incubated for 2 hours at room temperature with the appropriate secondary antibodies diluted in PBS-TB. Following secondary antibody incubation, the sections were rinsed for 30 minutes in PBS and coverslipped under 90% glycerol in PBS. Images were collected using a 20X objective on an Olympus BX51-WI microscope equipped with conventional fluorescence filters (Cy3:528–553 nm excitation, 590–650 nm emission; Cy2/Alexa 488: 460–500 nm excitation, 510–560 nm emission), a Hamamatsu ER, DVC high-speed camera, linear focus encoder, and a 3-axis motorized stage system interfaced with Neurolucida software (MBF Bioscience, Essex, VT, USA).

### In vivo electrophysiological recording of human peripheral sensory fibers

Single-unit axonal recordings (microneurography) were performed from the right posterior antebrachial cutaneous, radial, or superficial peroneal nerve of 41 healthy participants (19 males and 22 females; 19 to 41 years). Participants were comfortably seated in an adjustable chair with legs and arms stretched out (and hand pronated), supported by vacuum pillows, and covered in a blanket if they reported as feeling cold.

Under real-time ultrasound guidance (LOGIQ P9, GE Healthcare, Chicago, IL, USA), the target nerve was impaled with an insulated tungsten recording electrode (FHC Inc., Bowdoin, ME, USA). Adjacent to that, an uninsulated reference electrode was inserted just under the skin. A high-impedance preamplifier (MLT185 headstage) was attached to the skin near the recording electrode and used together with a low-noise high-gain amplifier (FE185 Neuro Amp EX, ADInstruments, Oxford, UK). Once the electrode tip was intrafascicular, single LTMRs were searched for by soft-brush stroking, and single HTMRs were searched for by coarse-brush stroking, pinching, and hair tugging in the fascicular innervation zone while making minute electrode adjustments.

All recorded afferents were mechanically responsive and divided into subtypes based on established criteria^68,112,113^. Mechanical threshold and receptive field size were determined using Semmes-Weinstein monofilaments (nylon fiber; Aesthesio, Bioseb, Pinellas Park, FL, USA). Mechanical threshold was defined as the weakest monofilament to which the unit responded to in at least 50% of trials. Hair deflection was tested with a small pair of forceps, carefully avoiding skin contact while manipulating the hair. Further, force measurements were performed to ensure that no skin/hair pulling occurred. Conduction velocity of the recorded afferent was estimated from latency responses to surface electrical stimulation of the receptive field (FE180 Stimulus Isolator, ADInstruments, Oxford, UK). Electrically and mechanically evoked spikes were compared on an expanded time scale to confirm they originated from the same unit. Thermal responsiveness was tested by placing a Peltier probe (7.4 x 12.2 mm, T09, QST.Lab, Strasbourg, France) onto the receptive field. After recording baseline activity for at least 30 s (with the thermode in contact with the receptive field) at a neutral temperature of 30°C, a series of cooling (down to 0°C) and warming (up to 50°C) stimuli were delivered at 30-s intervals. If needed, the thermode was mounted on a stand for better stability.

To test *TRPV1* expression, capsaicin (Capsina 0.075%, Bioglan AB, Malmö, Sweden) was topically applied to the receptive field. After 1 minute, the skin was wiped clean, and the emergence of any spontaneous spiking activity from the recorded afferent was monitored. *TRPM8* expression was tested by placing an ethanol-soaked gauze pad (90% ethanol as control) onto the receptive field followed by menthol solution (400 mg of 40% L-menthol dissolved in 90% ethanol, Sigma-Aldrich, Inc., Schnelldorf, Germany^114^). The gauze pad was covered with an adhesive film to prevent the evaporation of ethanol. After 5 minutes, the skin was wiped clean and the emergence of any spontaneous spiking activity from the recorded afferent was monitored. During these procedures, we documented the participants’ verbal descriptions of what they felt, and if there was no obvious sensation, the procedure was repeated.

Neural activity was sampled at 20 kHz and recorded using the ADInstruments data acquisition system (LabChart software v8.1.24 and PowerLab 16/35 hardware PL3516/P, Oxford, UK), then exported to Spike2 (v10.13, Cambridge Electronic Design Ltd., Cambridge, UK). Recorded action potentials were carefully examined offline on an expanded time scale. Threshold crossing was used to distinguish action potentials from noise with a signal-to-noise ratio of at least 2:1, and spike morphology was confirmed by template matching. Recordings were discarded if multiple units were present or if spike amplitudes were not distinct from the noise, preventing secure action potential identification.

## Figure Generation Software

Figures were generated in Powerpoint (Microsoft Office) and GraphPad Prism (v9, GraphPad Software Inc. La Jolla, CA, USA). Some cartoons were made partially in BioRender (BioRender, 2022, RRID:SCR_018361).

## Data availability

The raw and processed datasets for the single-soma sequencing of hDRG neurons reported in this study will be deposited into Broad Institute Single cell portal (https://singlecell.broadinstitute.org/single_cell) once the manuscript is accepted for publication.

Macaque (Kupari) data is available at https://www.ncbi.nlm.nih.gov/geo/query/acc.cgi?acc=GSE165569

Mouse (Zeisel) DRG data is available at http://loom.linnarssonlab.org/clone/Mousebrain.org.level6/L6_Peripheral_sensory_neurons.loom.

Mouse (Sharma) DRG data is available at https://www.ncbi.nlm.nih.gov/geo/query/acc.cgi?acc=GSE139088

## Code availability

Any custom code will be deposited to Github once the manuscript is accepted for publication. All analyses are based on previously published code and software.

---

**Supplementary Figure 1.**
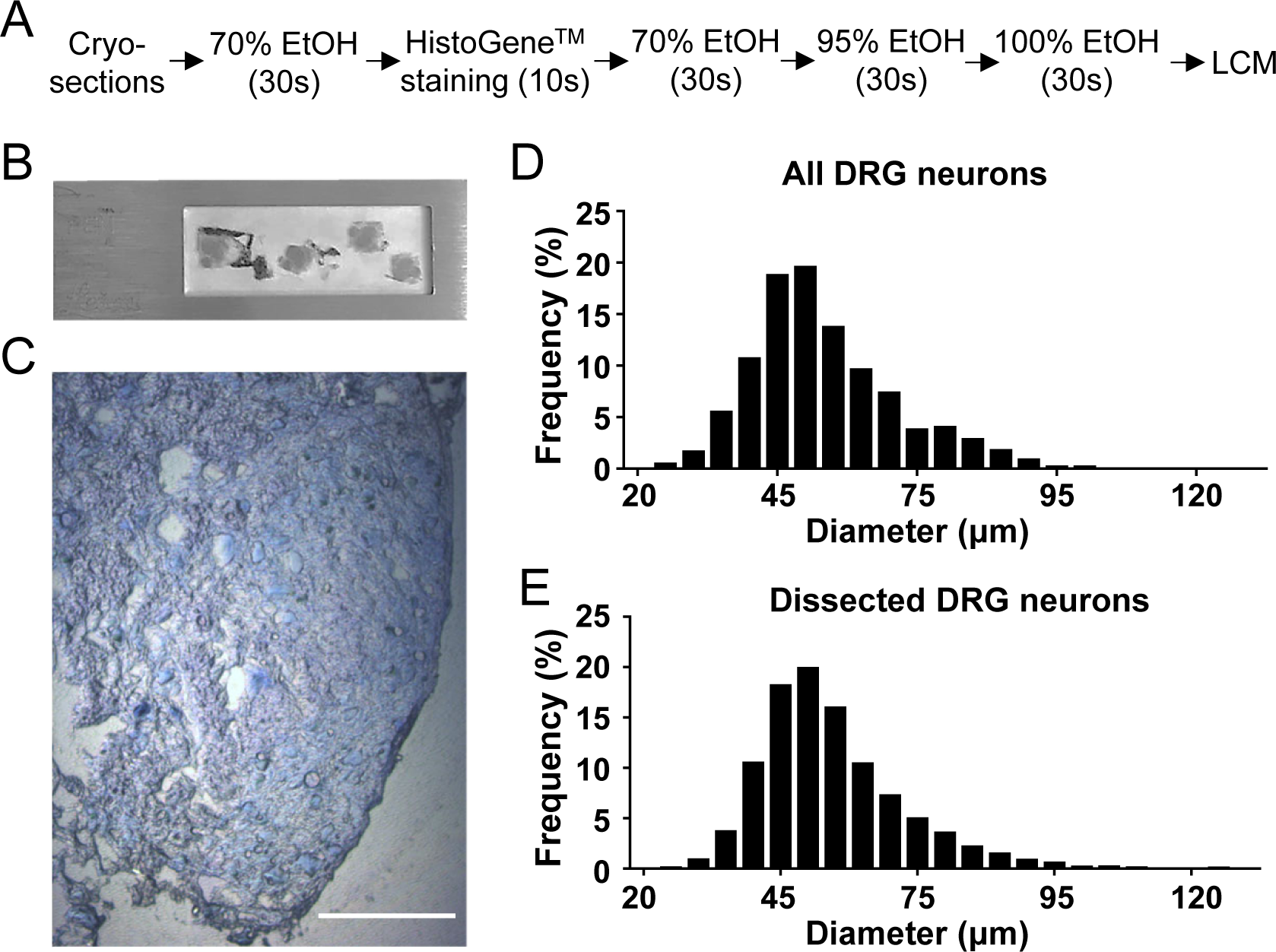

**Supplementary Figure 2.**
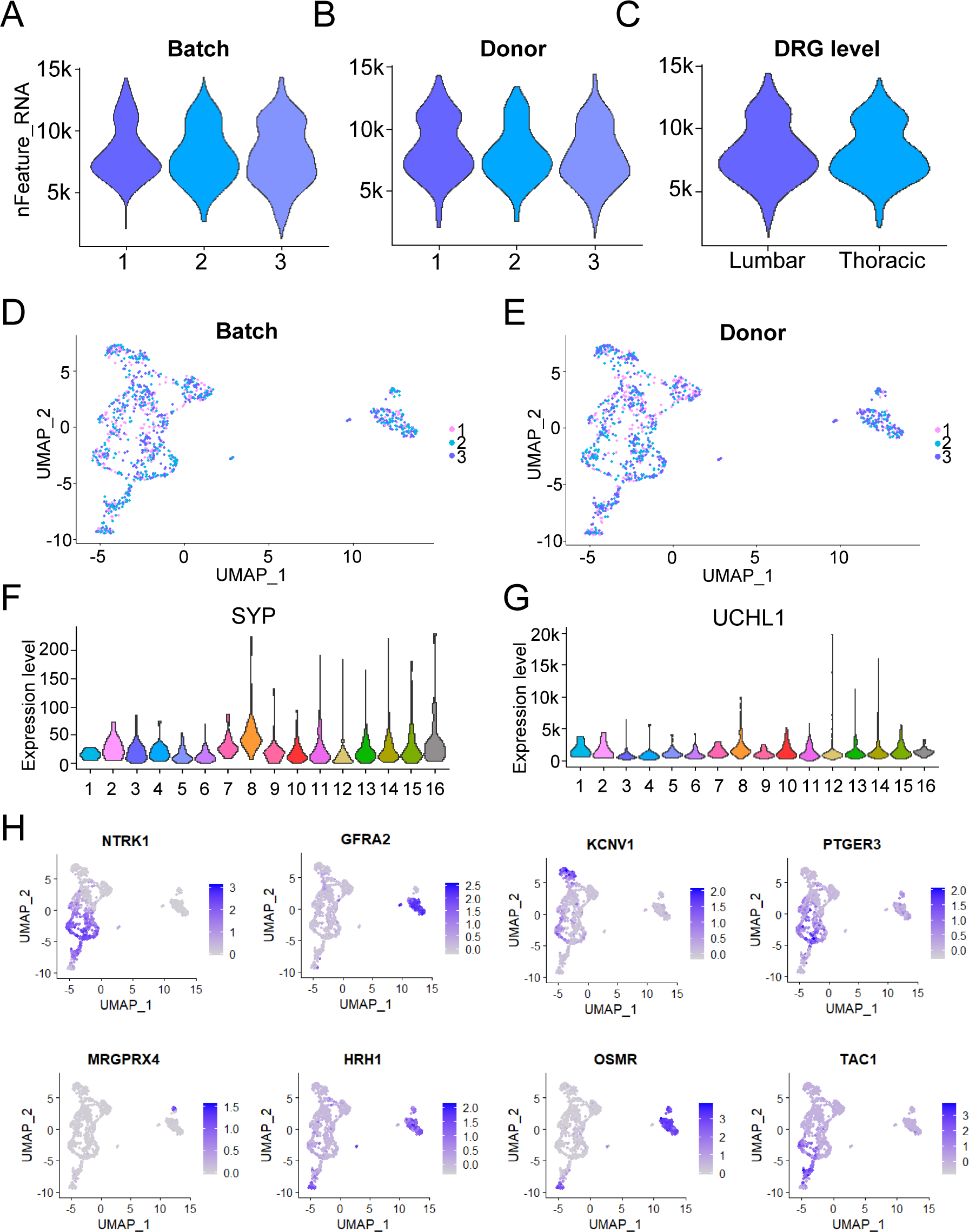

**Supplementary Figure 3.**
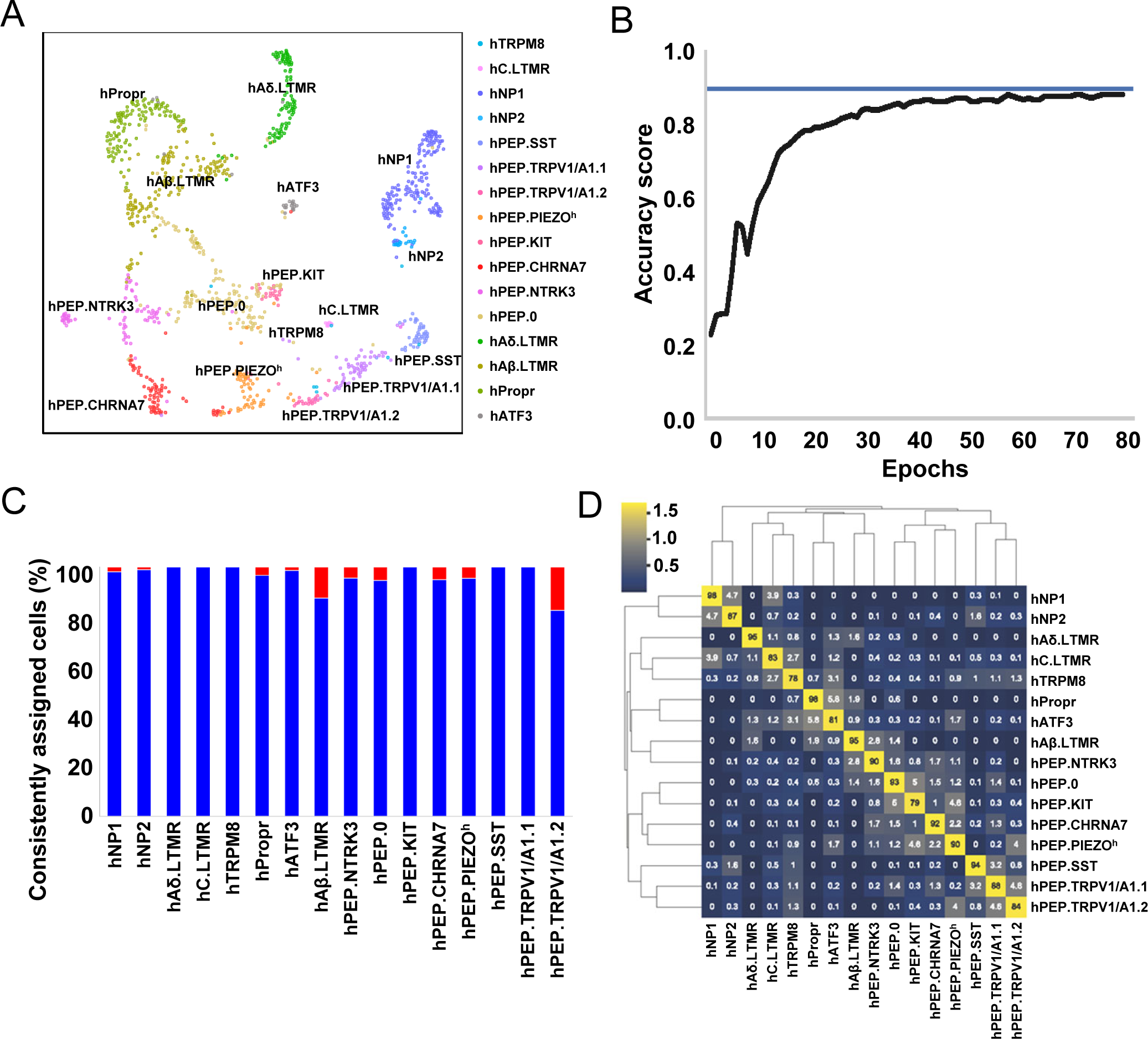

**Supplementary Figure 4.**
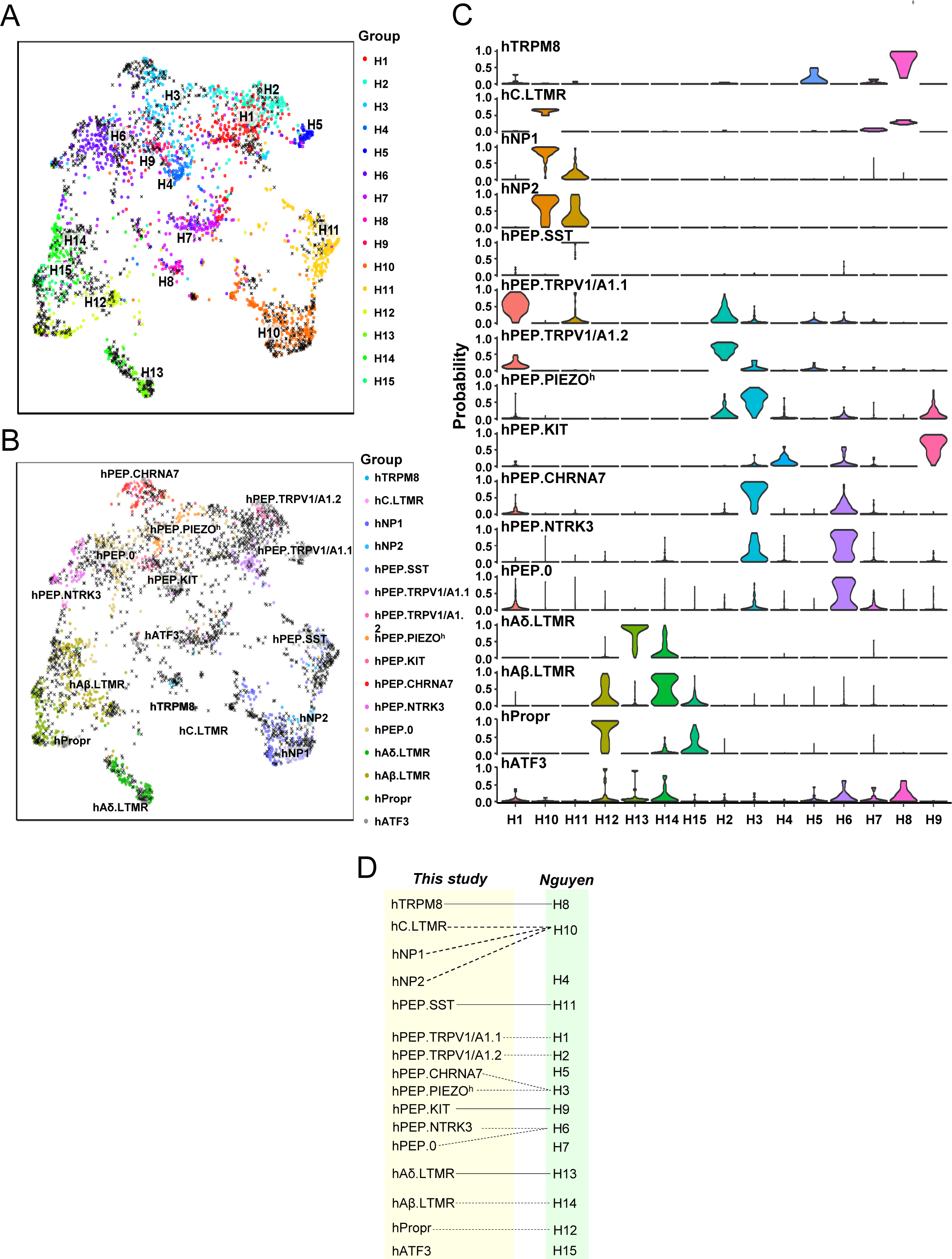

**Supplementary Figure 5.**
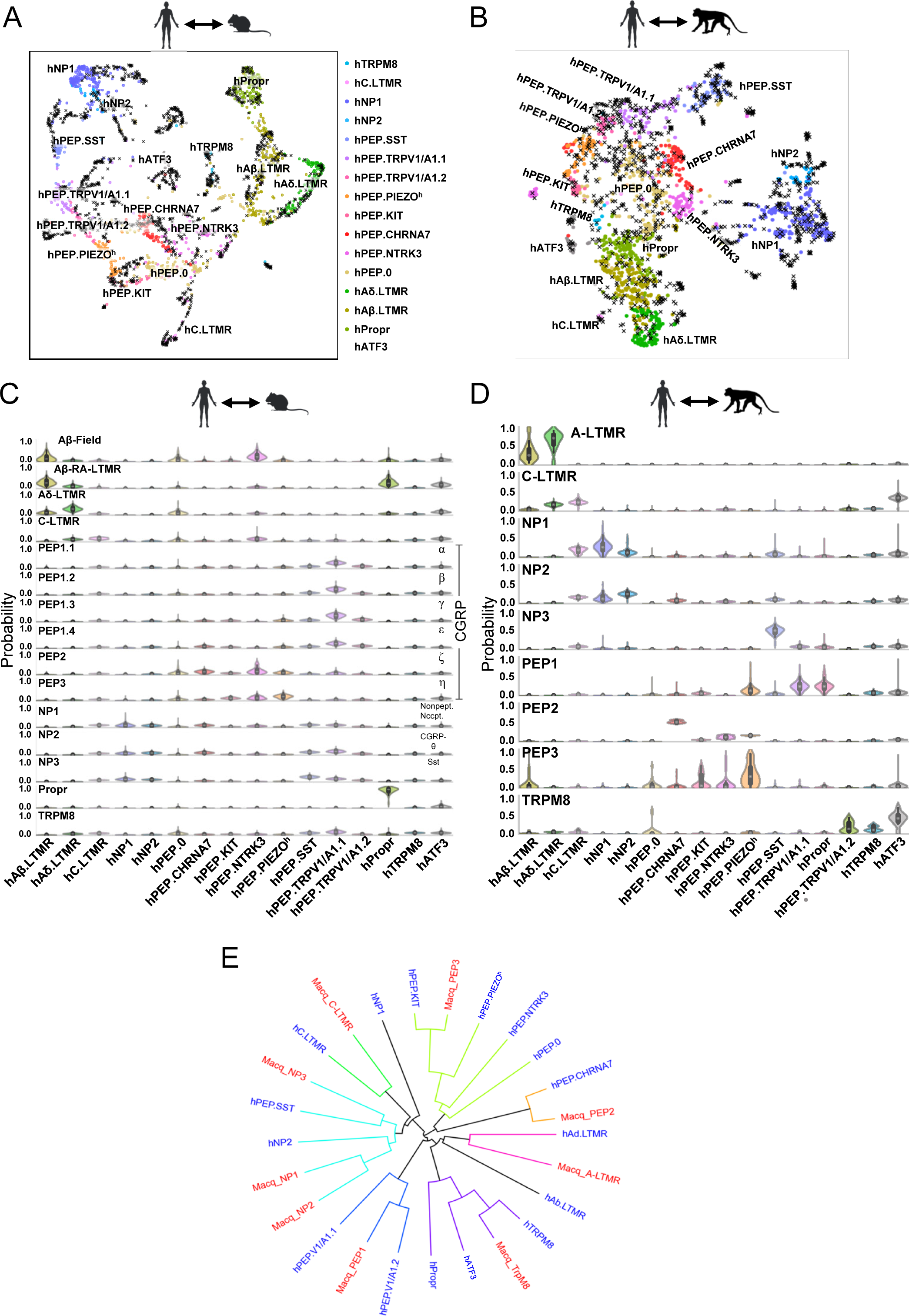

**Supplementary Figure 6.**
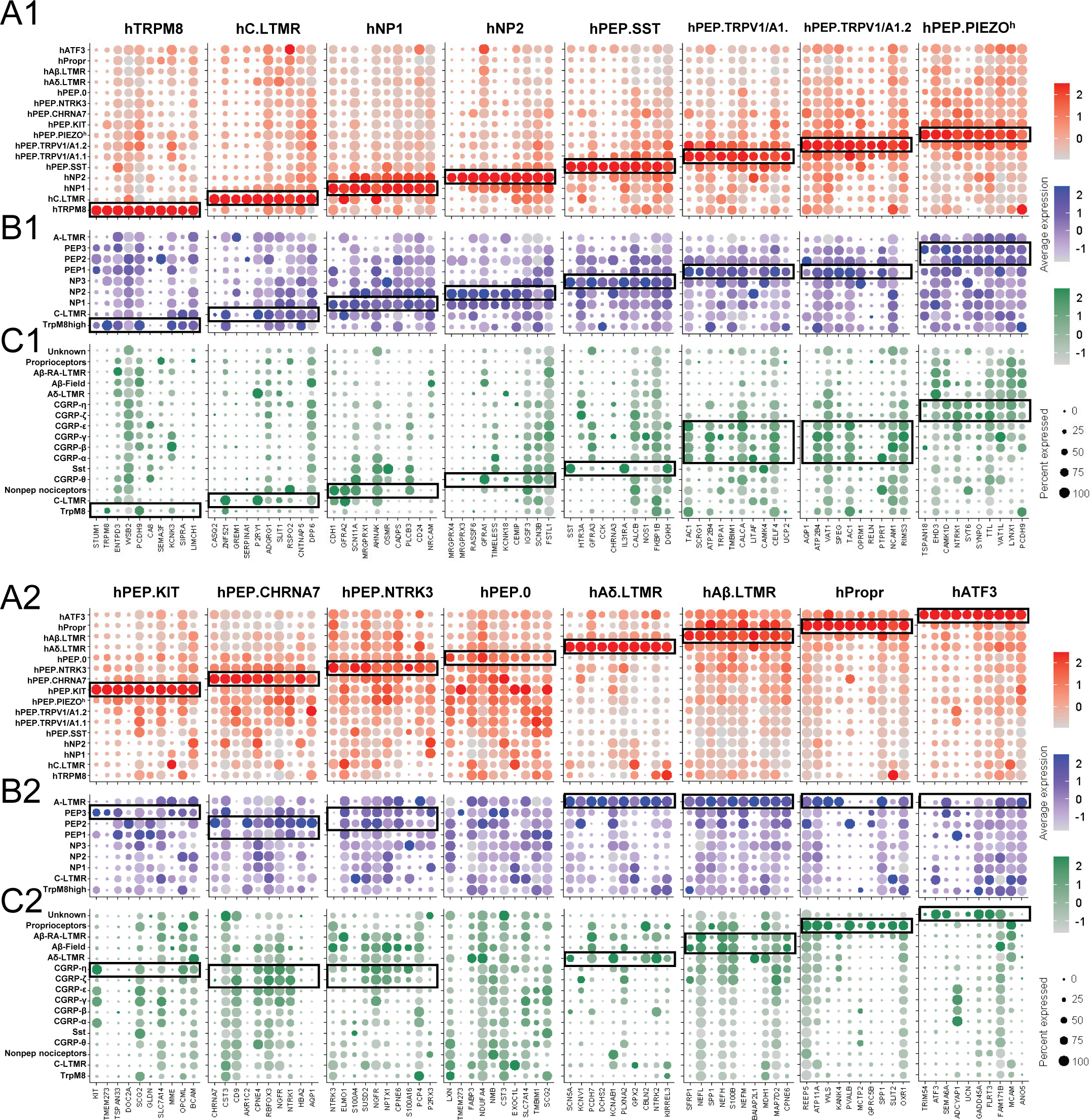

**Supplementary Figure 7.**
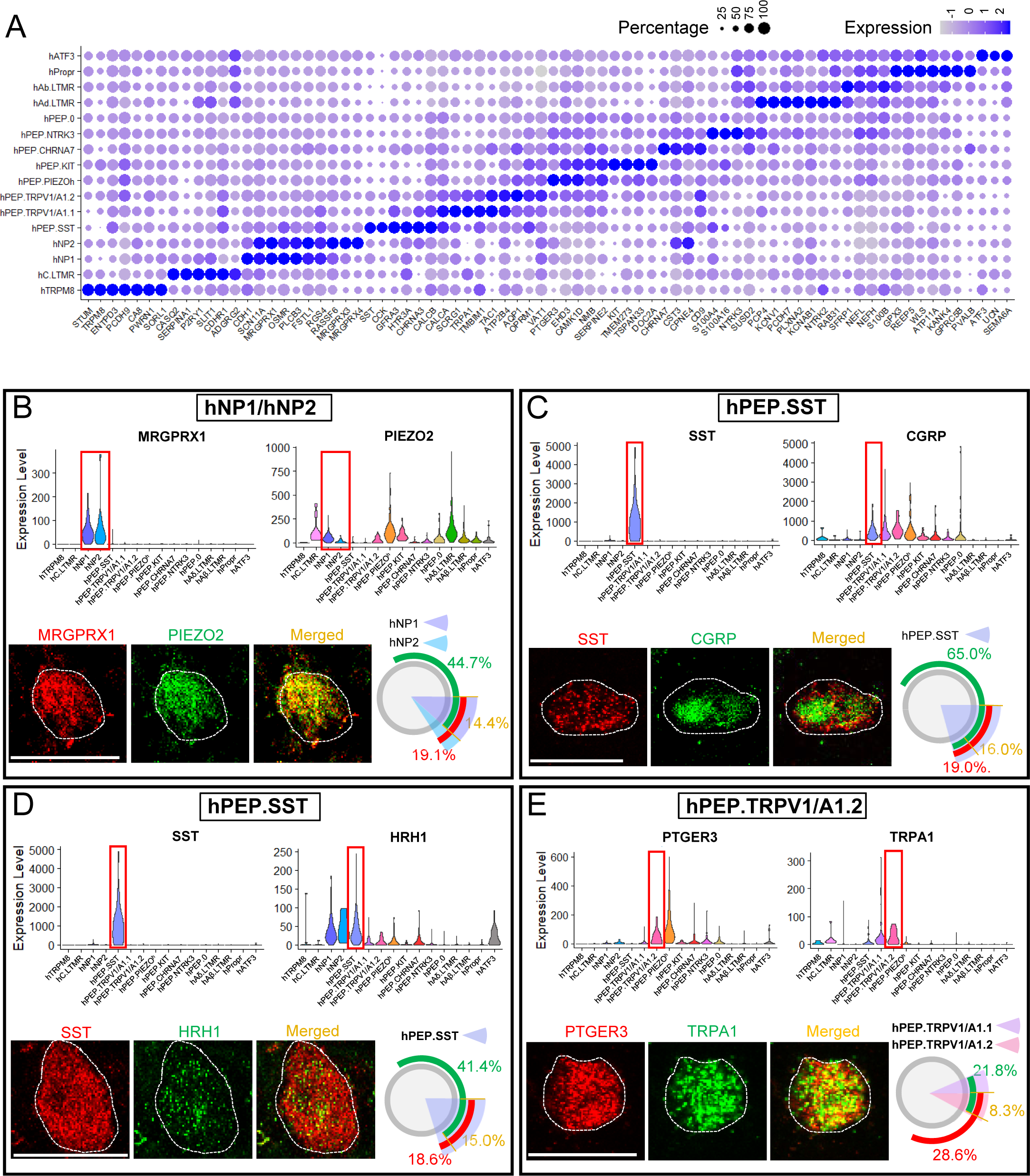

**Supplementary Figure 8.**
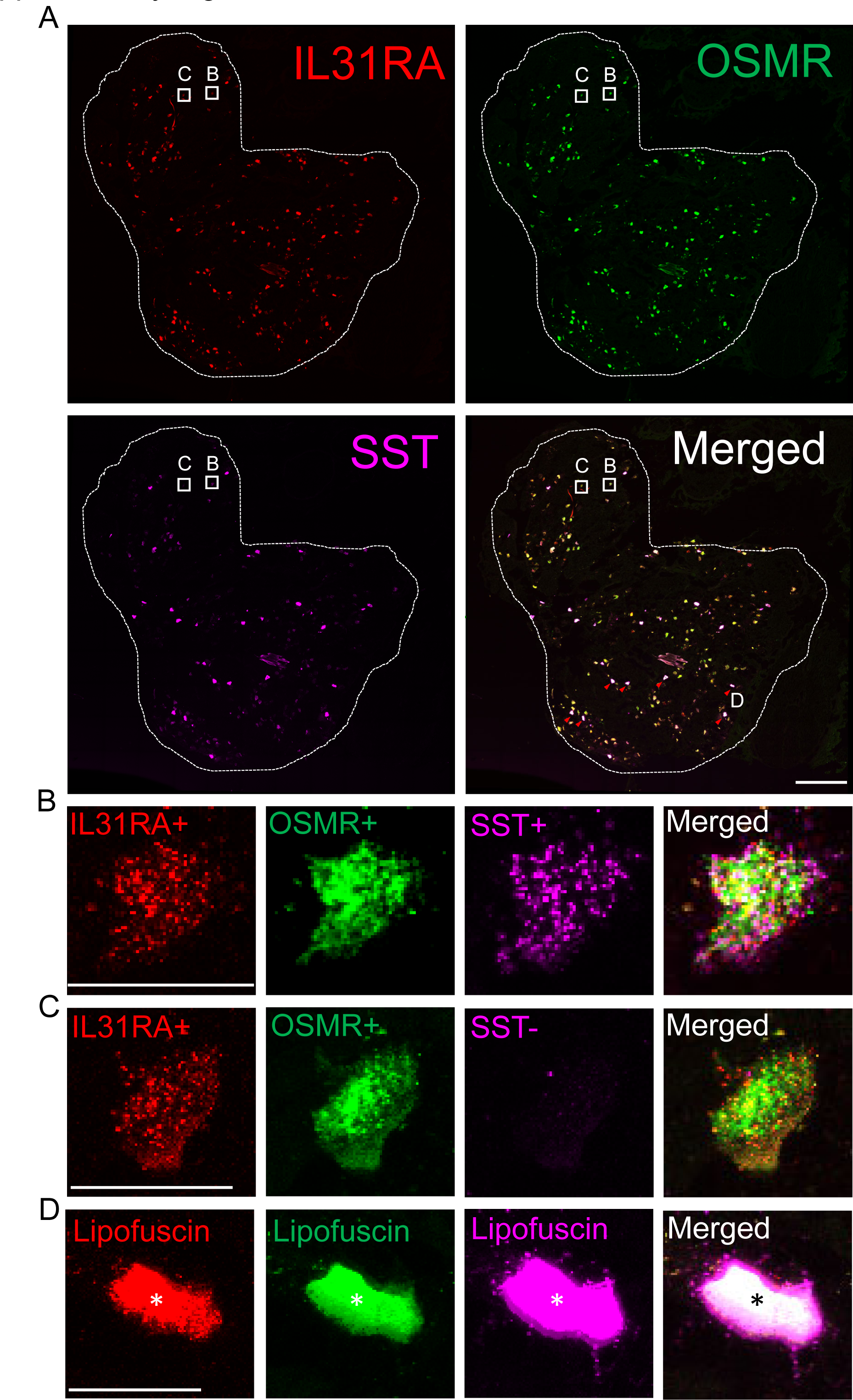

**Supplementary Figure 9.**
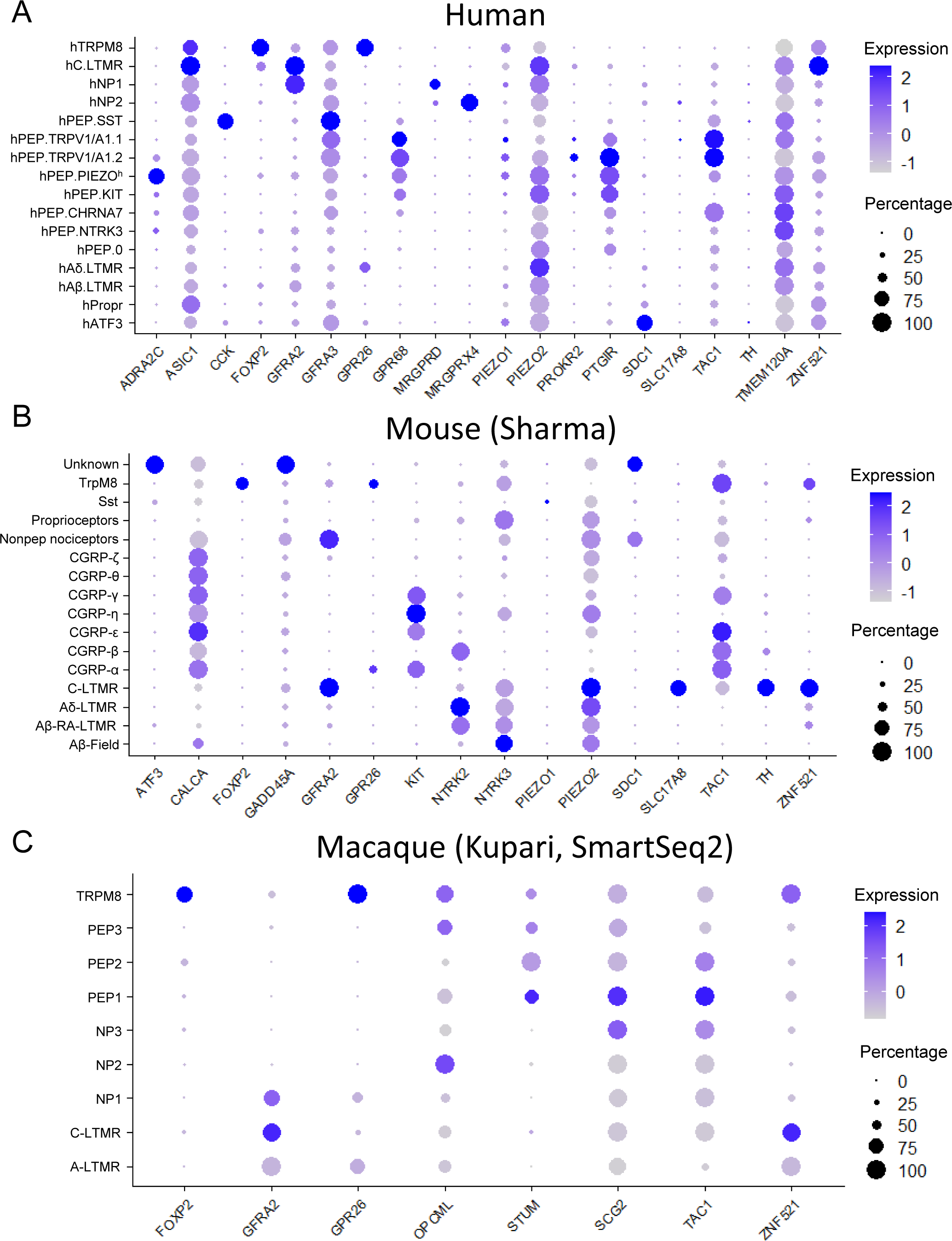

**Supplementary Figure 10.**
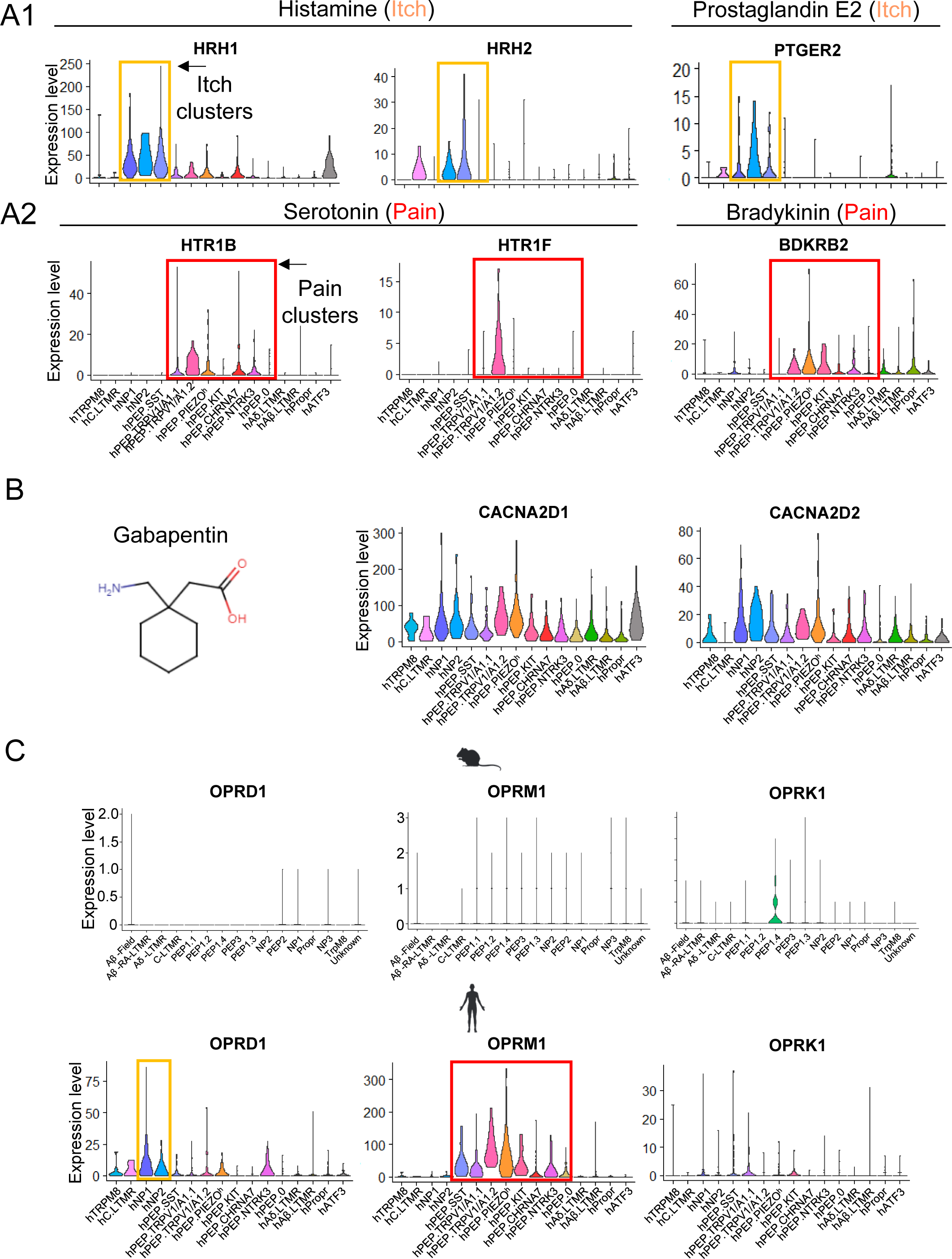

**Supplementary Figure 11.**
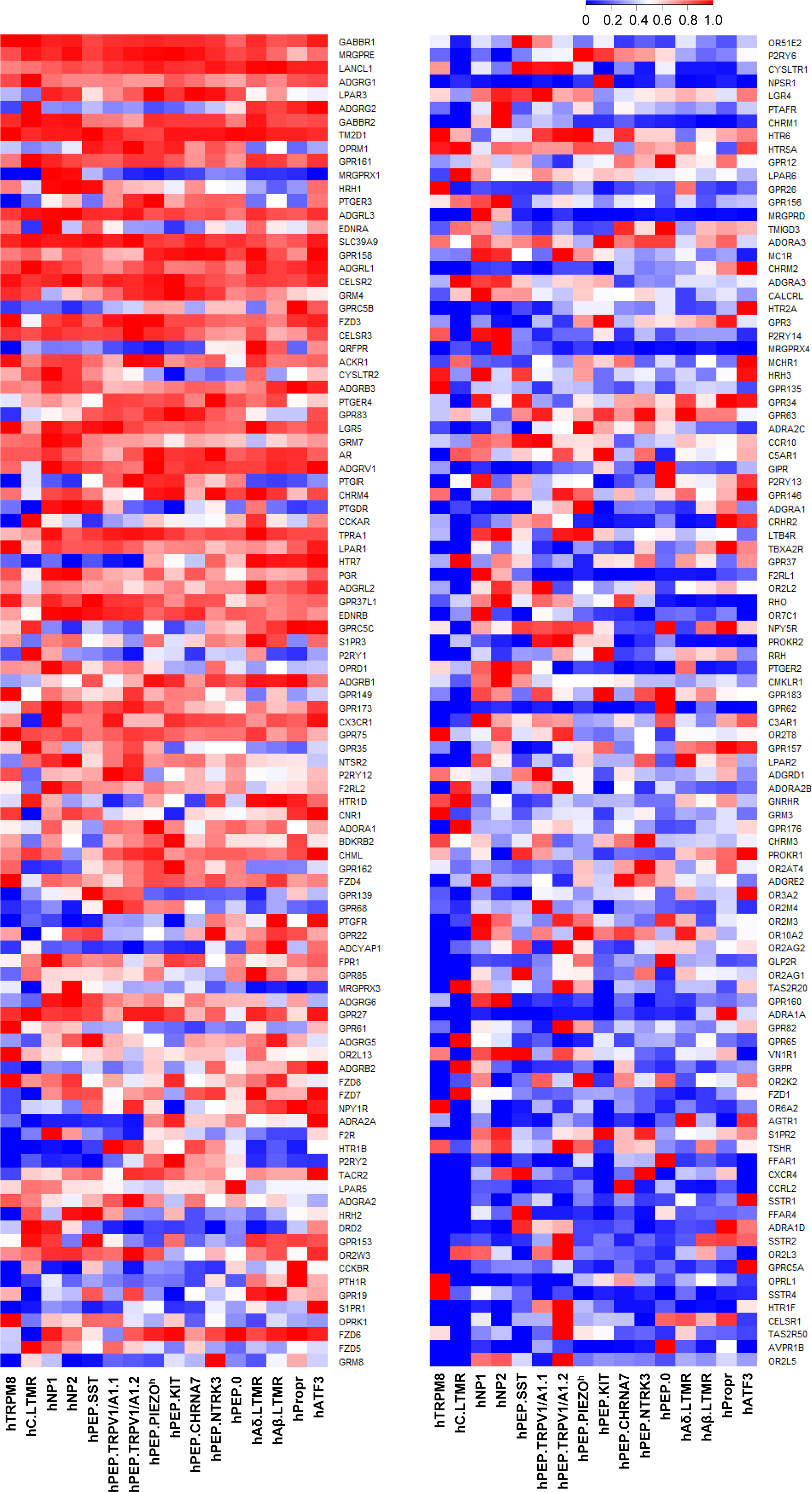

**Supplementary Figure 12.**
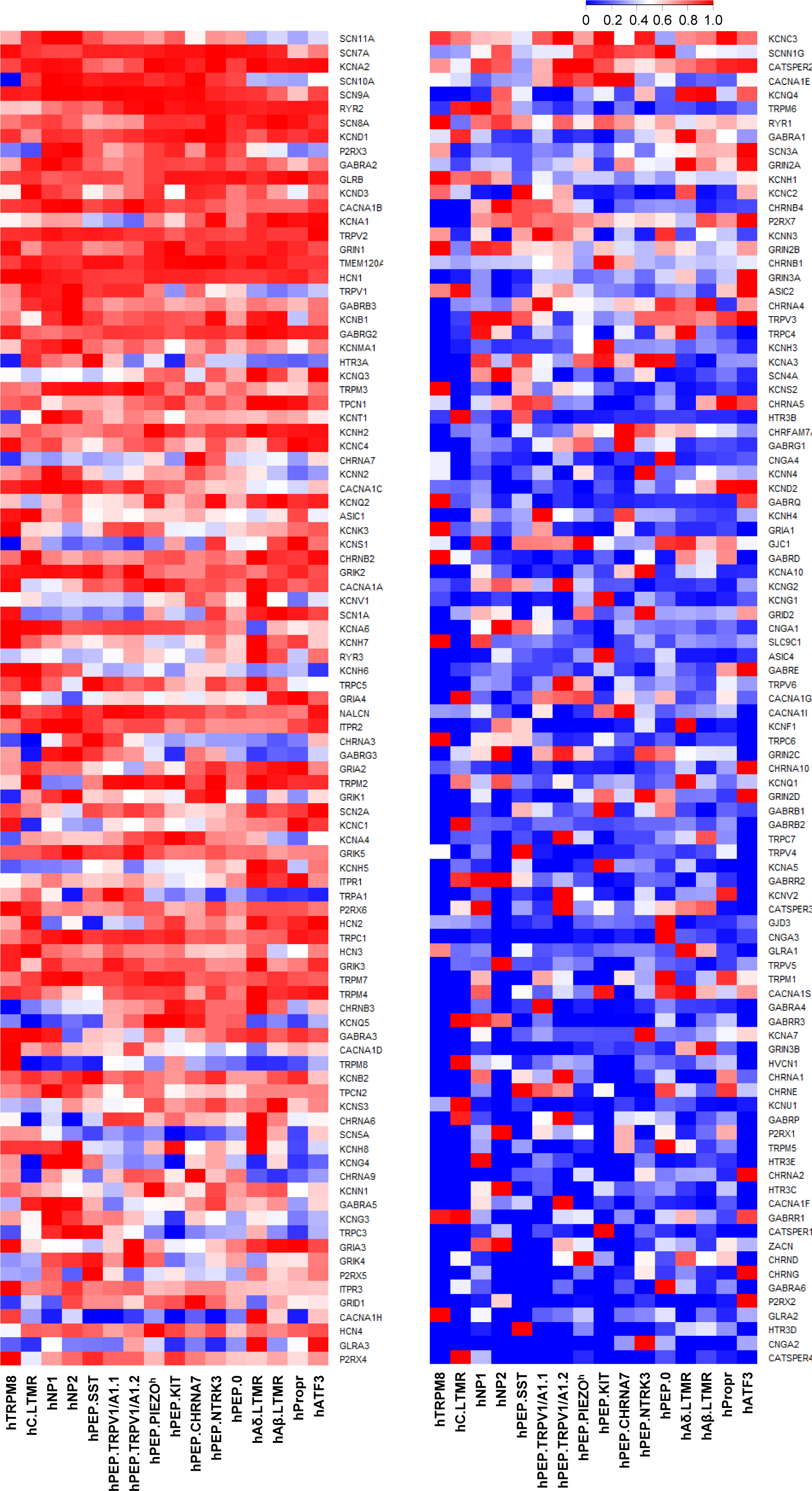

**Supplementary Figure 13.**
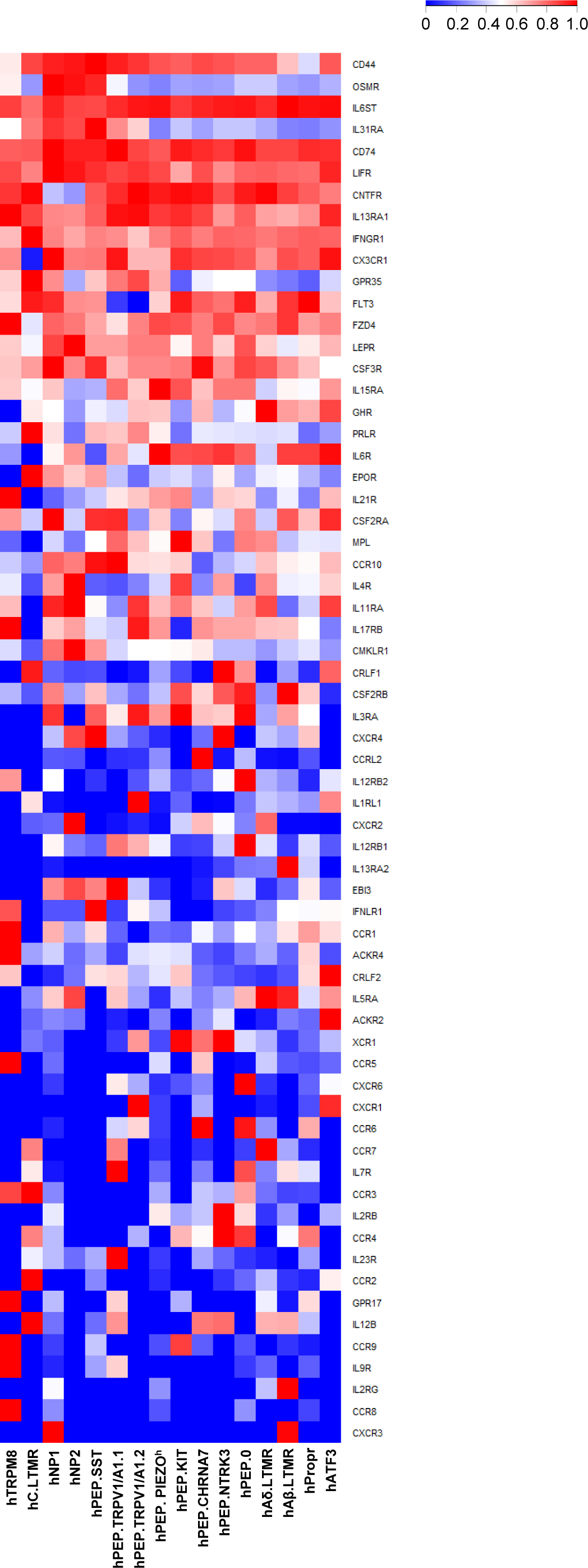

**Supplementary Figure 14.**
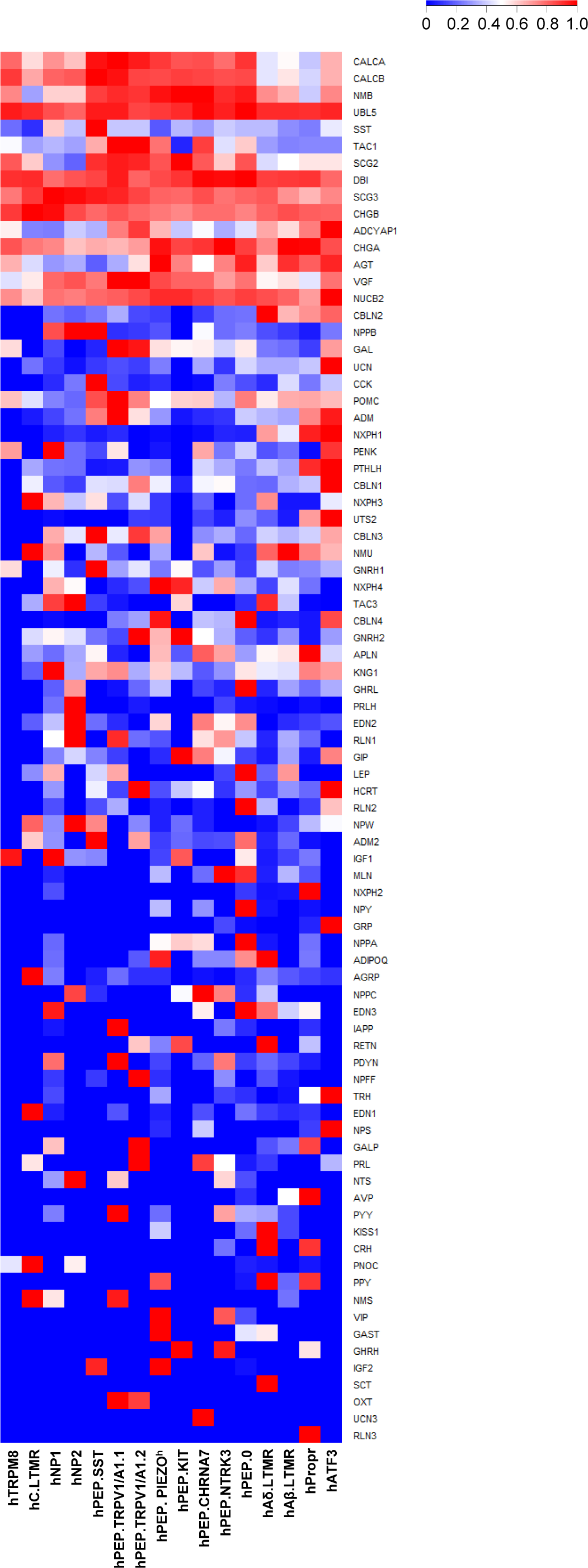

**Supplementary Figure 15.**
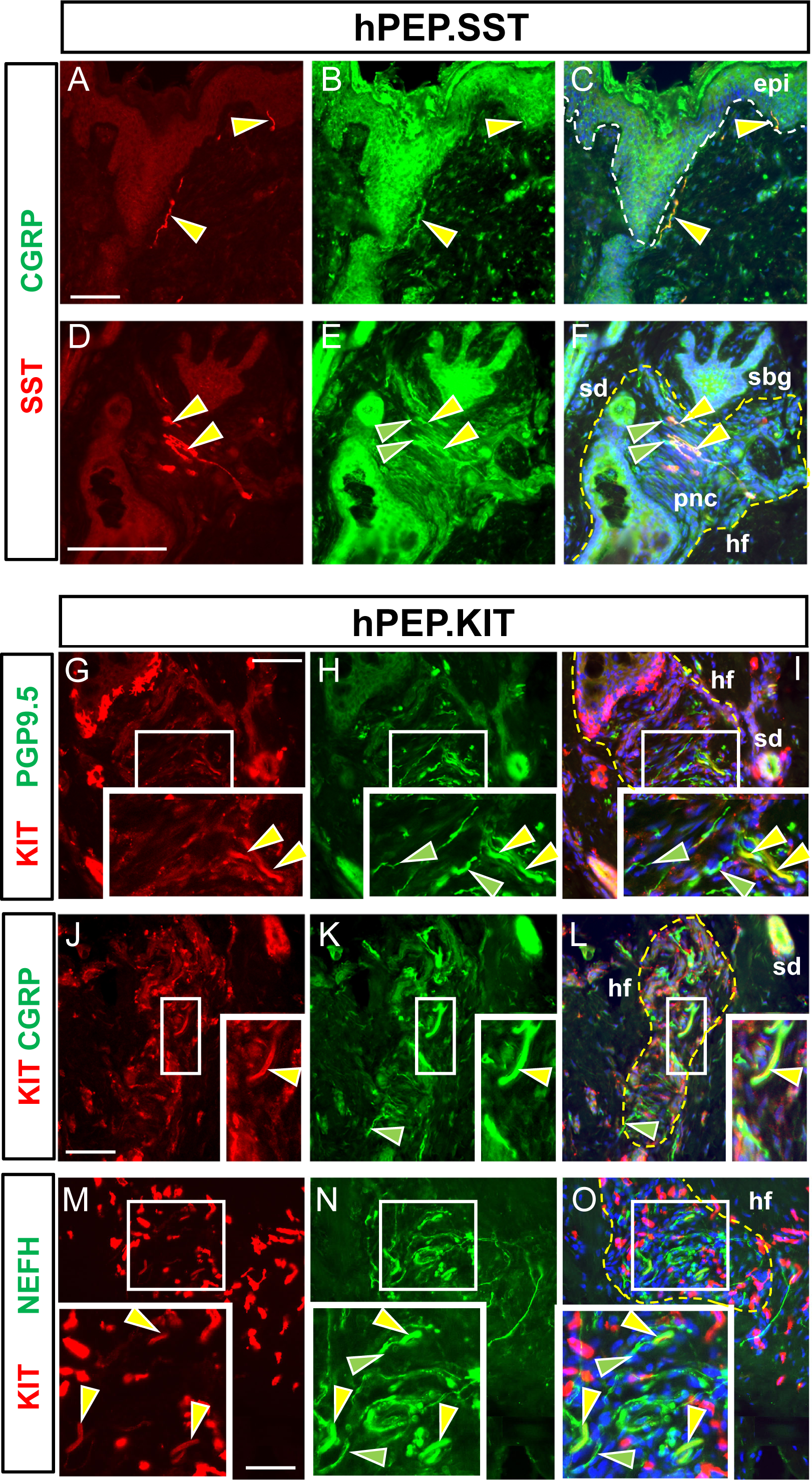

**Supplementary Figure 16.**
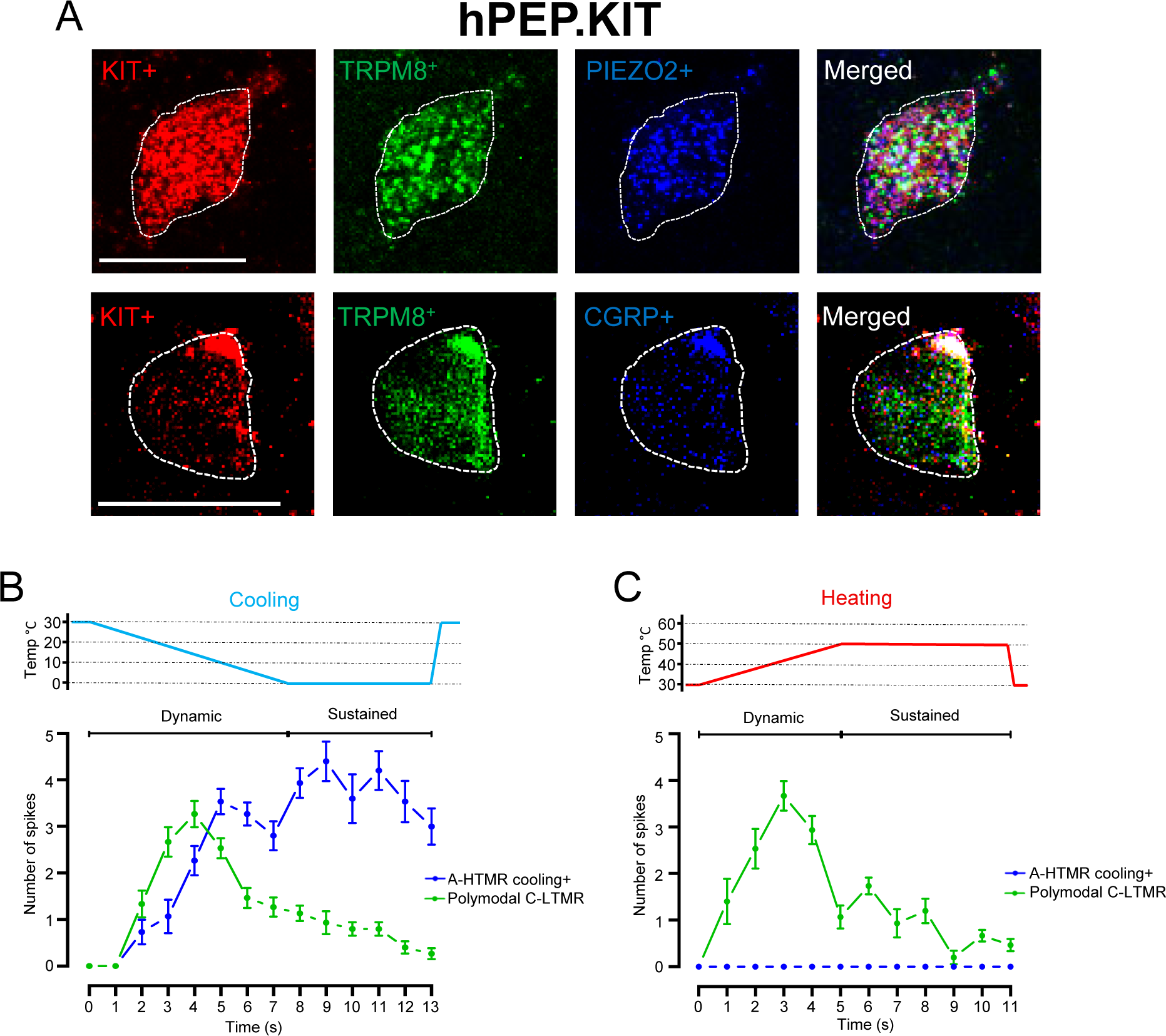

**Supplementary Figure 17.**
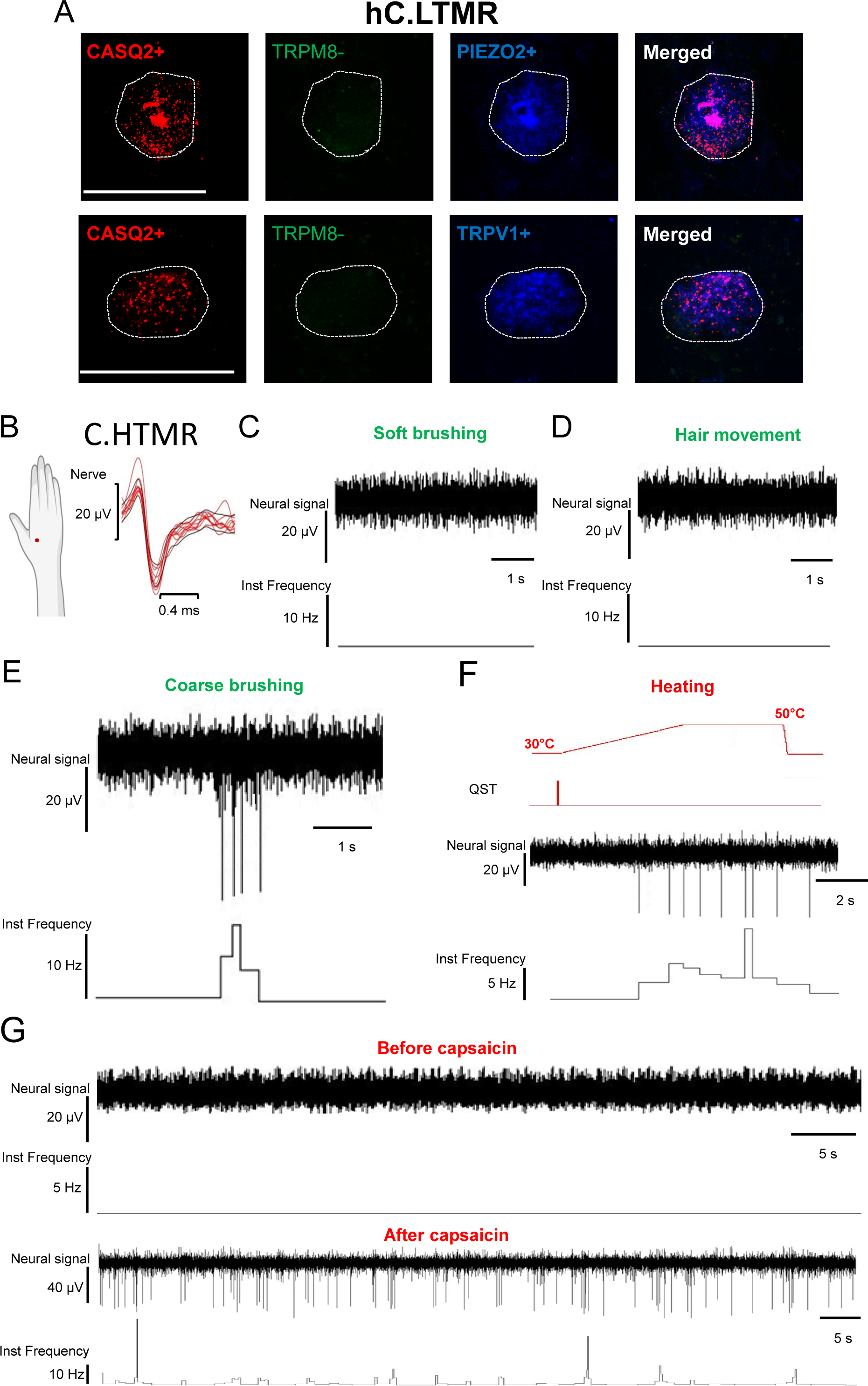

