## Supplementary table and figure legend for "Single-Soma Deep RNA Sequencing of Human Dorsal Root Ganglion Neurons Reveals Novel Molecular and Cellular Mechanisms Underlying Somatosensation"

**Supplementary table 1. Summary of donor information for DRG samples**

| **Tissue ID** | **DRG level** | **Donor ID** | **Age** | **Sex** | **Race** | **BMI** | **Date of sampling** |
| --- | --- | --- | --- | --- | --- | --- | --- |
| N2-RL5 | Lumbar 5 | 2012-08042 | 61 | female | Caucasian | 31.09 | 12/26/2020 |
| N2-RT12 | Thoracic 12 |  |  |  |  |  |  |
| N3-RL2 | Lumbar 2 | 2103-02139 | 56 | female | Caucasian | 32.44 | 03/19/2021 |
| N3-RT11 | Thoracic 11 |  |  |  |  |  |  |
| N4-RL3 | Lumbar 3 | 2106-05090 | 23 | male | Caucasian | 30.87 | 06/27/2021 |
| N4-RT12 | Thoracic 12 |  |  |  |  |  |  |

**Supplementary table 2. Donor screening** **criterion for human DRG samples**

| Age Range | 18-65 |
| --- | --- |
| Race | Any |
| Sex | No Preference |
| Required Disease(s) | Normal |
| Required Medication(s) | None |
| Required Surgeries(s) | None |
| Infectious Disease Testing Required | Yes |
| Sepsis | No |
| Increased Risk | Some Acceptable If Infectious Disease Negative |
| Acceptable Warm Ischemic Time (WIT) | Less than 60 |
| Acceptable Downtime | Less than 60 |
| Vent Time | Always Acceptable |
| Cancer | Never Acceptable |
| Chemotherapy | Never Acceptable |
| Radiation | Never Acceptable |
| Unacceptable Diseases | Diabetes - Type I (T1D, T1DM)  Diabetes - Type II (T2D, T2DM)  Diabetes |
| Acceptable Diseases | None |
| Unacceptable Surgeries | None |
| Acceptable Surgeries | None |
| Unacceptable Medications | None |
| Acceptable Medications | None |
| Tobacco Use | Always Acceptable |
| Alcohol Use | Some Acceptable  Quantity: 2 Drinks per Day  Day Duration: No Restrictions  Use Category: Moderate Drinking  Time Since last use: No Restrictions |
| Illicit Drug Use | Never Acceptable |

**Supplementary table 3. Summary of donor information for human skin samples**

| **Tissue ID** | **DRG level** | **Donor ID** | **Age** | **Sex** | **Race** | **Date of sampling** |
| --- | --- | --- | --- | --- | --- | --- |
| 868-01 | Left Ankle | CNGB-0868 | 60 | female | Caucasian | 03/27/2023 |
| 868-02 | Left Thigh |  |  |  |  |  |
| 868-03 | Left Hand |  |  |  |  |  |
| 868-04 | Right Ankle |  |  |  |  |  |
| 868-05 | Right Thigh |  |  |  |  |  |
| 868-06 | Right Hand |  |  |  |  |  |
| 892-01 | Left Hand | CNGB-0892 | 26 | female | Caucasian | 05/15/2023 |
| 892-02 | Left Thigh |  |  |  |  |  |
| 892-03 | Left Ankle |  |  |  |  |  |
| 892-04 | Right Hand |  |  |  |  |  |
| 892-05 | Right Thigh |  |  |  |  |  |
| 892-06 | Right Ankle |  |  |  |  |  |
| 793-01 | Left Hand | CNGB-0793 | 24 | male | Caucasian | 07/14/2023 |
| 793-02 | Right Hand |  |  |  |  |  |
| 793-03 | Right Thigh |  |  |  |  |  |
| 793-04 | Left Thigh |  |  |  |  |  |
| 793-05 | Left Ankle 1 |  |  |  |  |  |
| 793-06 | Left Ankle 2 |  |  |  |  |  |

**Supplementary figure 1. Isolation of hDRG neuronal soma by LCM**

(**A**) A workflow showing steps for brief staining of hDRG section before LCM. **(B)** An image showing the hDRG cryosections mounted on the LCM slide after brief staining. (**C**) A representative image showing the stained hDRG section. Scale bar, 400 μm (**D-E**) Size distribution of all DRG neurons (D) and dissected neuronal soma (E).

**Supplementary figure 2. LCM RNA-seq statistics, clustering, and additional marker gene expression**

(**A-C**) Violin plots showing total number of detected genes in different batches (A), donors (B), and DRG levels (C). (**D-E**) UMAP plots showing the contribution of individual batches (D) and donors (E) to each cluster. (**F-G**) Violin plots showing the expression of pan-neuronal markers *SYP* and *UCHL1*. (**H**) UMAPs showing expression pattern of additional canonical marker genes in each cluster.

**Supplementary figure 3. Validation of hDRG neuron clustering**

(**A**) Clustering of hDRG neurons by Conos. (**B**) The accuracy of neural-network classifier in learning hDRG neurons was visualized as learning curve. The Y-axis represents the learning accuracy, and the X-axis represents the training epoch numbers. Following the training epochs, the maximum accuracy plateaus of this learning curve reaching ~88%. (**C**) Percentage bar chart visualization, the consistency of assigned hDRG neurons by the neural-network scoring module. Red indicating the ratio of cells that were not consistently assigned to their defined cell-types, and blue indicating the ratio of consistently assigned cells. (**D**) hierarchical heatmap visualization of the probabilistic similarity across cell types, color from dark blue to yellow indicating the similarity score from low to high.

**Supplementary figure 4. Co-clustering with single-nucleus RNA-seq of hDRG neuron**

(**A-B**) Co-clustering of human single-soma RNA-seq and human single-nucleus RNA-seq (Nguyen). (**C**) Label transfer from human single-soma RNA-seq to human single-nucleus RNA-seq showing the cell type correlation between the two datasets. (**D**) Summary of correspondence in clusters between single-soma sequencing and single-nucleus sequencing (Nguyen). Solid lines depict clear match, dashed lines represent partial similarity.

**Supplementary figure 5. Cross-species comparison of sensory neuron types among human, macaque, and mouse**

(**A-B**) UMAPs of Conos co-clustering of human and mouse (A) or macaque (B) DRG neurons. These co-clustering were used for label propagation inferences shown on Fig. 2A & 2B. Each pair UMAPs were plotted with the same coordinates, the human clusters are shown in color, the small black uncolored crosses depict mouse and macaque clusters. (**C-D**) Probabilistic neural network probability scores of mouse (Sharma dataset) (C) and macaque (Kupari, SmartSeq2 dataset) (D) DRG neuron types tested on human trained module. (**E**) Hierarchical clustering of cell types in human and macaque.

**Supplementary figure 6. Comparison of marker gene expression across species**

Dot plots showing top ten specific marker genes selected from each hDRG neuron clusters expressed in human (**A1, A2**), macaque (Kupari) (**B1, B2**) and mouse (Sharma) (**C1, C2**) DRG neuron datasets. The black boxes highlighted the corresponding cell types based on the label transfer and neural-network scoring analysis.

**Supplementary figure 7 Additional marker genes and their expression validation in hDRG neurons**

(**A**) Dot plot showing the top marker genes expressed in each hDRG neuron cell types. (**B-E**) Marker genes for specific labeling of each cluster and validation by multiplex FISH for hNP1 and hNP2 (B), hPEP.SST (C-D), and hPEP.TRPV1/A1.2 (E) The fluorescent images show the detected transcripts in one example hDRG neuron (cell body outlined by the white dashed line). Circle charts next to images show quantification: the arcs indicate the percentage of neurons positive for the given marker gene over all sampled DRG neurons. The sector shaded areas indicate the approximate percentage of each cell type over the total quantified hDRG neurons. N=2, B (188 neurons total), C (100 neurons total), D (220 neurons total), E (192 neurons total). Scale bar, 50 μm.

**Supplementary figure 8. Examples to show multiplex FISH of hDRG neuron and the non-specific lipofuscin autofluorescence**

(**A**) Low magnification Images of entire hDRG sections after performing multiplex FISH using probes for *IL31RA* (red), *OSMR* (green), *SST* (purple), and the merged. (**B-C**) The white box areas (B and C) on merged image in (A) are shown at higher magnification. An example of IL31RA, OSMR and SST triple positive neuron shown in (B) and an example of IL31RA and OSMR positive but SST negative neuron shown in (C). (**D**) An example of lipofuscin autofluorescence (red arrowhead in the merged image in (A)) indicated by asterisk (*) showed non-specific signals, which are excluded for analysis. Scale bar, 500 μm in (A), 50 μm in (B-D)

**Supplementary figure 9. Expression of additional genes expressed in human, mouse and macaque DRG neuron cell types**

Dot plots showing the expression of additional genes in human, mouse and macaque DRG neuron cell types.

**Supplementary figure 10. Potential anti-itch and anti-pain drug targets expression in hDRG neurons**

(**A1-A2**) Violin plot showing the receptor expression of itch and pain inducing chemicals in hDRG neurons. (**B**) Violin plot showing the expression of potential gabapentin targets in hDRG neurons. (**C**) Violin plot showing the expression of opioid receptors in mouse and human DRG neurons.

**Supplementary figure 11. Expression of GPCRs in hDRG neurons**

Heatmap showing the expression of GPCRs in hDRG neurons. Genes were ranked by average expression level in all DRG neurons.

**Supplementary figure 12. Expression of ion channels in hDRG neurons**

Heatmap showing the expression of ion channels in hDRG neurons. Genes were ranked by average expression level in all DRG neurons.

**Supplementary figure 13. Expression of chemokine receptors in hDRG neurons**

Heatmap showing the expression of chemokine receptors in hDRG neurons. Genes were ranked by average expression level in all DRG neurons.

**Supplementary figure 14. Expression of peptides in hDRG neurons**

Heatmap showing the expression of peptides in hDRG neurons. Genes were ranked by average expression level in all DRG neurons.

**Supplementary figure 15. Immunostaining of different types of sensory fibers in the human skin using molecular makers identified by single-soma deep RNA-seq dataset**

(**A-F**) Double immunostaining of human skin sections showing SST+ sensory fibers are a subset of CGRP+ sensory fibers in the dermis, the dermis-epidermis junction, and entering the epidermis (A-C) and near the hair follicle (D-F). The white dashed line indicates the dermis-epidermis junction. Yellow arrows indicate SST+/CGRP+ sensory afferents, green arrows indicate CGRP+ only sensory afferents (**G-O**). Double immunostaining of KIT with PGP9.5 (G-I), NEFH (J-L) and CGRP (M-O) in human hairy skin showing KIT+ sensory afferents are a subset of CGRP+/NEFH+ sensory afferents around the hair follicles. Yellow dashed lines outline hair follicles. Yellow arrows indicate double positive sensory afferents, green arrows indicate PGP9.5+ only or CGRP+ only or NEFT+ only sensory afferents. Epi, epidermis; hf, hair follicle; sd, sweat gland duct; pnc, pilo-neural complex; sbg, sebaceous gland. Scale bar, 100 μm.

**Supplementary figure 16. Expression of *TRPM8, PIEZO2* and *CALCA* in the hPEP.KIT population, and cooling and heating response properties of A-HTMR and C-LTMR units.**

(**A**) Validation of the expression of *TRPM8*, *PIEZO2* and *CGRP* in the hPEP.KIT population by multiplex FISH. (**B & C**) Responses of A-HTMR cooling+ and polymodal hC.LTMR units to temperature changes. Each data point represents the mean (± SEM) responses of 5 A-HTMR cooling+ or 5 polymodal hC.LTMR units, tested in triplicate. For hC.LTMR responses, adjustments were made for conduction delay based on the latency of electrically triggered spiking for those recorded afferents. (B) Number of action potentials evoked during dynamic (30°C to 0°C at 4°C per second) and sustained (0°C for 5.5 seconds) phases of cooling. (C) Number of action potentials evoked during dynamic (up to 50°C at 4°C per second) and sustained (~50°C for 6 seconds) phases of heating.

**Supplementary figure 17. Expression of *TRPM8, PIEZO2 and TRPV1* in hC.LTMR population and recordings from a C-fiber mechano-heat nociceptor**

(**A**) Validation of the expression of *TRPM8*, *PIEZO2* and *TRPV1* in hC.LTMR by multiplex FISH. (**B**) Spike activity of a C-fiber mechano-heat nociceptor in response to repeated stimulations of the receptive field, superimposed on an expanded time scale. (**C-G**) Responses of a C-fiber mechano-heat nociceptor to soft brushing (C), hair movement (D), coarse brushing (E), and heating (F) and capsaicin (G). Conduction delay was adjusted based on the latency of electrically triggered spiking for that recorded afferent. Note, different scaling.
